# Comparative 3D ultrastructure of *Plasmodium falciparum* gametocytes

**DOI:** 10.1101/2023.03.10.531920

**Authors:** Felix Evers, Rona Roverts, Cas Boshoven, Mariska Kea-te Lindert, Julie M.J. Verhoef, Robert E. Sinden, Anat Akiva, Taco W.A. Kooij

## Abstract

Despite the enormous significance of malaria parasites for global health, some basic features of their ultrastructure remain obscure. In this study, we apply high-resolution volumetric electron microscopy to examine and compare the ultrastructure of *Plasmodium falciparum* gametocytes of both genders and in different stages of development as well as the more intensively studied asexual blood stages revisiting previously described phenomena in 3D. In doing so, we challenge the widely accepted notion of a single mitochondrion by demonstrating the presence of multiple mitochondria in gametocytes. We also provide evidence for a gametocyte-specific cytostome variant. Furthermore, we generate, among other organelles, the first 3D reconstructions of endoplasmic reticulum (ER), Golgi apparatus, and extraparasitic structures in gametocytes. Assessing interconnectivity between organelles, we find frequent structural appositions between the nucleus, mitochondria, and apicoplast. We provide evidence that the ER is a promiscuous interactor with numerous organelles and the trilaminar membrane of the gametocyte. Public availability of these volumetric electron microscopy resources of wild-type asexual and sexual blood-stage malaria parasites will facilitate reinterrogation of this global dataset with different research questions and expertise. Taken together, we reconstruct the 3D ultrastructure of *P. falciparum* gametocytes in high detail and shed light on the unique organellar biology of these deadly parasites.

## Introduction

Parasites from the genus *Plasmodium* are the causative agents of malaria. This mosquito-borne infectious disease remains a huge burden on global public health with more than 200 million cases and >600 thousand fatalities in 2021 alone^1, 2^. Humanity’s efforts to combat this disease have shown impressive results during the years 2000 – 2015 but have since stalled. This is largely driven by continuing emergence of resistance to all frontline antimalarials^3^ as well as the inability of most antimalarials to combat directly the asymptomatic but transmissible sexual stages^4^. To stop parasite transmission and eliminate malaria, there is an urgent need for drugs with novel mechanisms of action, particularly those that are effective against sexual stages. The apicomplexan phylum, among which are the malaria parasites, diverged very early in eukaryotic evolution and some basic cellular processes and structures are comparatively poorly understood despite their immense significance for global health. In particular, *Plasmodium* organelle biology has been shown to deviate from standard eukaryotic models and varies drastically between the different life-cycle stages. Furthermore, Apicomplexa are almost exclusively parasitic, which drives additional divergence in cellular features related to their survival strategy, such as feeding, locomotion, secretion, invasion, and adaptation to their respective hosts. As a result, we are in the situation where there is urgency to find drug or vaccine candidates for cellular systems that are relatively poorly understood. In the case of *Plasmodium*, this holds particularly true for life-cycle stages, like the sexual gametocyte stages, that are less accessible than the pathogenic asexual blood-stage parasites (ABS).

Classical electron microscopy studies in the period from 1965-2000 provided key insights into the ultrastructure of malaria parasites and laid the groundwork for much of our understanding of sexual-stage biology^*5*–*8*^. There have been a few noteworthy serial sectioning electron microscopy (ssEM) applications^9–12^, however, these early studies lacked the advanced quantitative, volumetric, and computational methods available today and did not have the context of our current molecular understanding of malaria parasite biology. With the advent of novel and, more importantly, increasingly accessible high-resolution volumetric approaches such as focused ion beam milling - scanning electron microscopy (FIB-SEM), serial block-face scanning electron microscopy (SBF-SEM), array tomography (AT), and expansion microscopy (ExM), we have made great advances in recent years. Features of ABS and oocysts and their respective replication strategies have been investigated via FIB-SEM with great success^13, 14^. ssEM and AT have recently been applied to elucidate the reputed role of nuclear microtubules in driving the characteristic elongation of developing gametocytes^15^ and general measurements of gametocytes and insights underpinning the inner membrane complex (IMC) have been gained using SBF-SEM^16^. ExM has been used to elucidate microtubule dynamics across gametocyte development and during activation, and immunofluorescence-based studies have demonstrated that the mitochondrion dramatically branches and enlarges in the gametocyte stages^17–19^. Additionally, electron micrographs have shown that in contrast to the acristate mitochondrion in ABS, the gametocyte mitochondrion is highly cristate^6, 20, 21^. Yet, other organelles, such as the endoplasmic reticulum (ER), the Golgi apparatus (Golgi), and the cytostome, and their interconnectivity have not been subject to specific microscopic investigations in gametocytes and our general understanding of their morphology and putative interactions is largely derived from model eukaryotes or extrapolated from *Plasmodium* ABS.

In this study, we investigate the ultrastructure of gametocytes using high resolution FIB-SEM and ssSEM volumetric imaging. We compare our ultrastructural gametocyte findings with our ABS data and other high-resolution volumetric studies. In doing so, we find evidence for a morphologically distinct cytostome in gametocytes, multiple mitochondria per cell, indications for widespread inter-organelle interactions, generate high-resolution renderings of various organelles and visualize 3D morphology and distribution of external structures in gametocytes. Additionally, we aim to create a reusable image resource for the malaria research community to investigate more cells from the FIB-SEM and ss-SEM data, contrast their own findings with a reference point, and reinterrogate the datasets with their specific biological question, perspective, and expertise.

## Results and Discussion

### General morphology of the gametocyte

The mature gametocytes show their typical crescent shape that is well known from light microscopy-based imaging (Fig. 1A-B, Movies S1-2). The surrounding red blood cell (RBC) also conformed to this general shape, with the Laveran’s bib, a narrow ridge spanning between the tips of the parasite crescent, being evident in all stage IV and V gametocytes (Fig. 2). We also identified contorted mature gametocytes with a twisted appearance including a narrowing of the cytoplasm, a frequent aberrant morphology known from Giemsa preparations of gametocyte cultures (Fig. S1). We were surprised to find that aside from the deviating morphology, no other signs of poor health or differences of internal structures could be recognized in these contorted gametocytes. Immature gametocyte stages (stage II-IV) were identified based on their well-characterized straighter or more compact appearance and incomplete inner membrane complex (Fig. S2). With increased maturity, the overall electron density of the RBC cytoplasm decreases and an electron lucent corona surrounding the gametocyte becomes apparent, indicative of hemoglobin depletion (Fig. S3). While we were unable to confidently assign sexes to earlier stages, we relied on differing patterns of hemozoin distribution together with the frequency and appearance of osmiophilic bodies (OBs) to differentiate male and female stage IV/V gametocytes. Whereas previously cytoplasmic density of ribosomes^22^ was widely used to discriminate genders, we could not detect such differences in our preparations. OBs are vesicles that are abundant in females and thought to play a role during gametocyte egress from the RBC. *P. falciparum* males have been suggested to contain fewer OBs or even lack them completely^5, 23^. Our data show that *P. falciparum* males, contain vesicles that share morphological characteristics with OBs (Fig. 1B, Fig. S4A, Movie S2). As in *Plasmodium berghei*, these structures are much fewer in number in males than in females^24^. Furthermore, all or almost all OBs in male gametocytes have a “tail” that occasionally interconnects the vesicles - which is found only in a small subset of OBs in females (Fig. S4). This tail has been observed previously in a thin section of a female *Plasmodium cathemerium* gametocyte and a *Plasmodium gallinaceum* gametocyte^25^ but is largely absent from previous microscopic data of *P. falciparum* gametocytes. It would be interesting to investigate further whether these different observations reflect differences in underlying biology of the analyzed parasites or methodological differences. One possible explanation is that paucity of prior observations might stem from a higher likelihood of identification in high-resolution volumetric data as opposed to individual thin slices or differing staining procedures that make these structures visible. In the rodent malaria parasite, *P. berghei*, male osmiophilic bodies have been shown to have a distinct but overlapping proteome compared to OBs identified in female gametocytes^24^. However, no morphological differences were evident from the presented representative micrographs. We hypothesize that the putative OBs identified in male *P. falciparum* gametocytes also differ in protein composition as males do not stain positive for the Pfg377 protein that readily stains OBs of female gametocytes^23^. Supporting this hypothesis a recent preprint suggests presence of two functionally distinct subtypes of vesicles with different proteomes that aid in gametocyte egress, one of which – in line with our data - does not show any evidence for gender specificity^26^.

**Figure 1.**
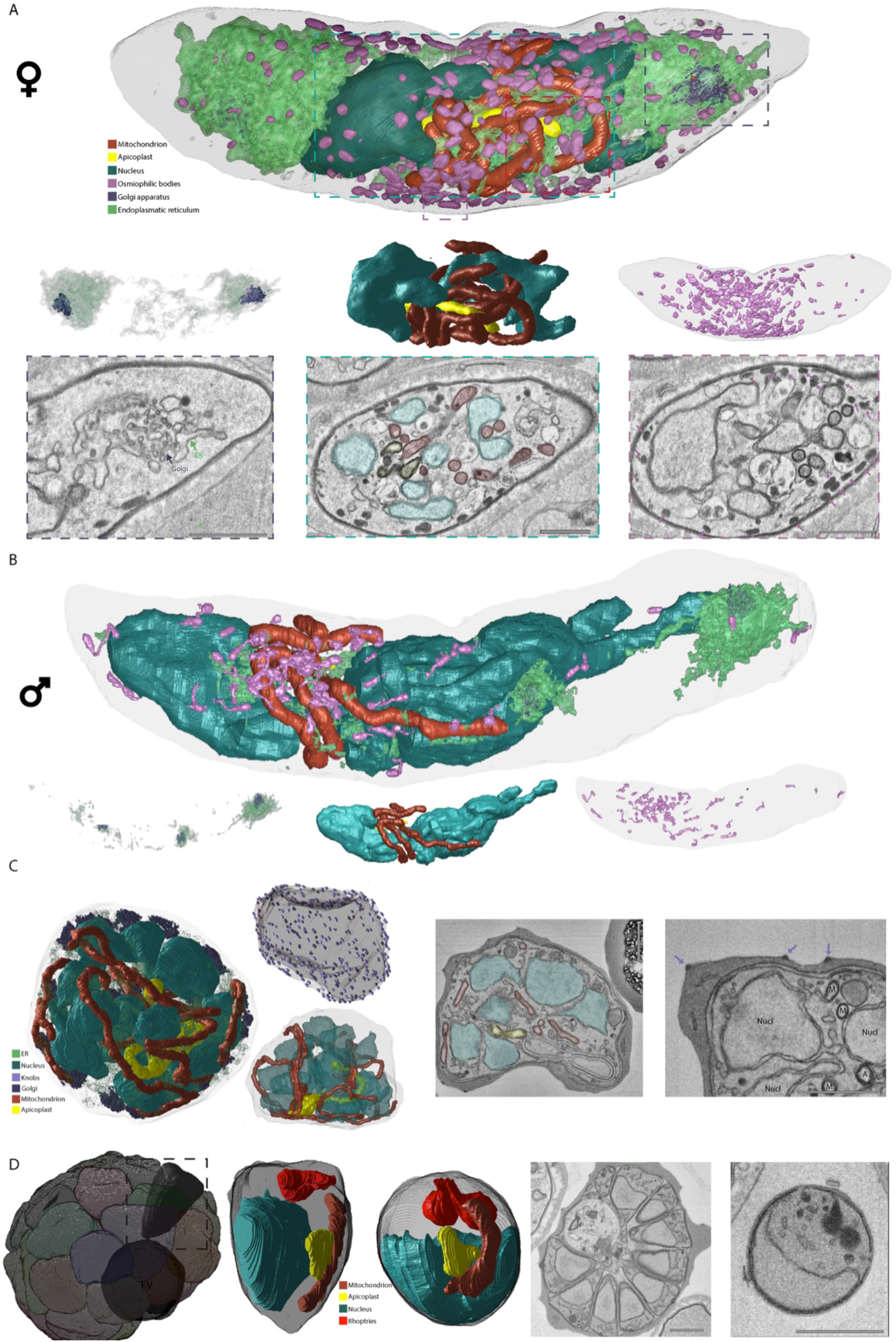
Ultrastructural features and renderings of mature gametocytes, schizont and segmented schizont. **A.** Rendering of ultrastructural features of a mature female gametocyte and specific renderings of Golgi and ER distribution (left panel), relation of nucleus/apicoplast/mitochondrion (middle panel) as well as distribution and appearance of osmiophilic bodies. Appearance of rendered structures in exemplary micrographs is matched to rendered features through color-coded dashed lines. Scale bars = 1 μm. **B.** Rendering of ultrastructural features of a mature male gametocyte and specific renderings as in (A). Note the relative paucity of ER, Golgi, and osmiophilic bodies relative to the female gametocyte. The osmiophilic bodies differ morphologically with overall thinner appearance and frequent tail-like extensions. **C.** Rendering of ultrastructural features of an early schizont and specific rendering of relation of nucleus/apicoplast/mitochondrion as well as individual knob distribution. Appearance of ultrastructural features is shown in two exemplary micrographs. Scale bars = 1 μm. M = Mitochondrion; ER = Endoplasmic reticulum; Nucl = Nucleus; FV = Food vacuole. **D.** Rendering of daughter merozoites within a segmented schizont and rendering of one intracellular and one extracellular merozoite with ultrastructural features. Overview on appearance of the two merozoite shapes is shown in two exemplary micrographs. Scale bars = 1 μm.

**Figure 2.**
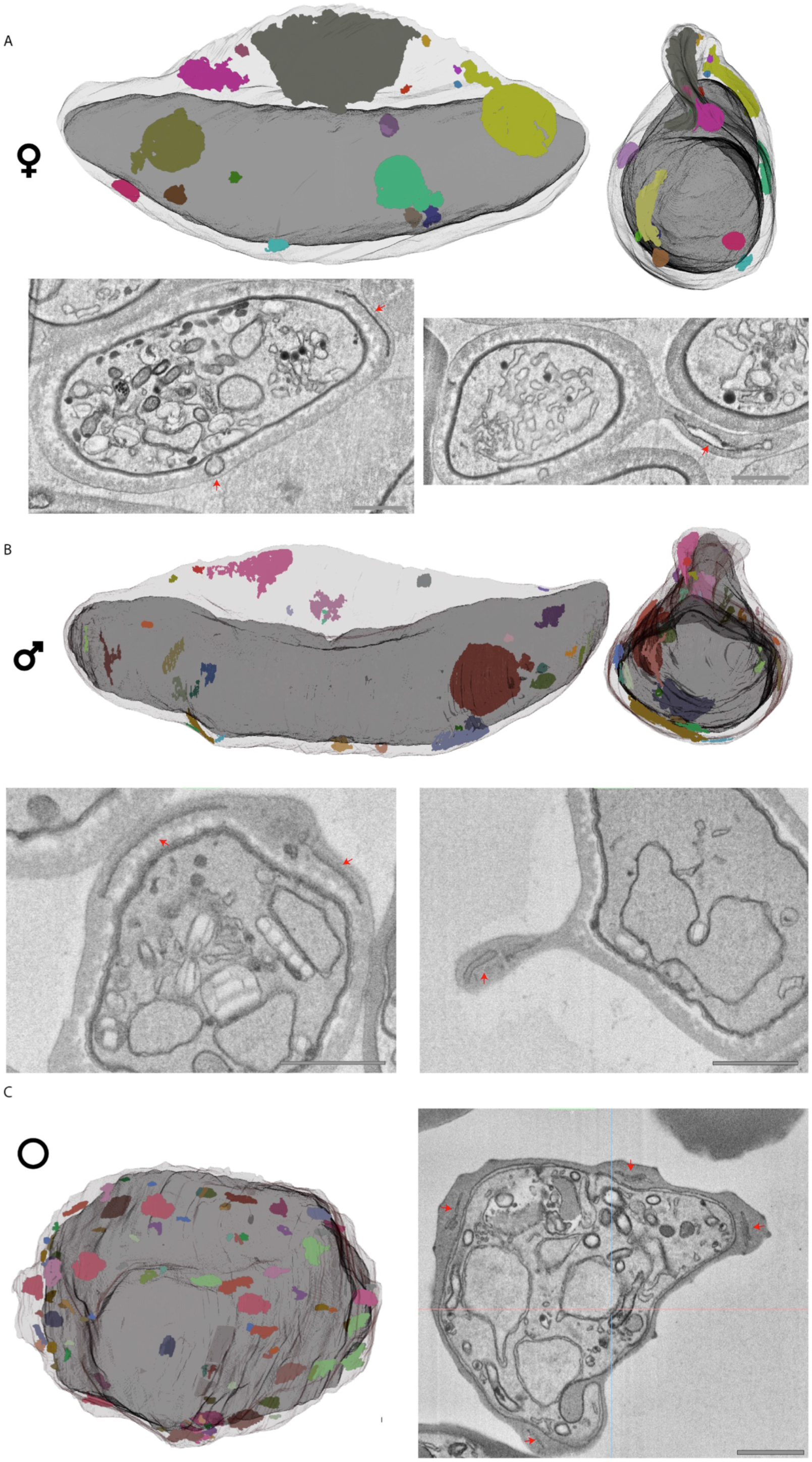
Extraparasitic structures differ in gametocytes and ABS. (**A**) Mature female and (**B**) male gametocyte, and (**C**) schizont rendered with RBC outline and extraparasitic structures in different color for each membrane structure. Gametocyte renderings are shown from two angles. Examples of extraparasitic structures in micrographs are highlighted with red arrows. Scale bars = 1 μm.

To compare our gametocyte reconstructions with the more extensively studied ABS^13^, we reconstructed two representative cells: a schizont that is still undergoing nuclear division and a fully segmented schizont, (Fig. 1C-D, Movie S3). Overall 3D morphology and appearance of the structures in individual slices match those observed by Rudlaff *et al*. ^13^ (publicly available from EMPIAR-10392). The schizont contains ten distinct nuclei and extensive mitochondrial and apicoplast networks that permeate the whole cell. The segmenter is comprised of 31 fully formed daughter merozoites, each containing mitochondrion, apicoplast, rhoptry pair, ER, micronemes, and nucleus, and an additional (32^nd^) merozoite that lacks a nucleus. Even at this late stage of development, without exception, each merozoite is still connected to the residual body that surrounds the food vacuole (Fig. 1C, Movie S3). This connectivity is highlighted by one merozoite that is connected to the residual body through an elongated neck/tube (Fig. S5A). Furthermore, the residual body is connected to the RBC cytoplasm via two continuous tubes, presumably facilitating continued sustenance to the daughter merozoites until rupture. Our data also contain occasional extracellular merozoites, most likely due to schizonts rupturing in the interval between magnetic separation and fixation. Comparing released extracellular to the intraerythrocytic merozoites, we find that volumes of the overall merozoite and individual organelles are very similar, supporting the notion that the corresponding segmented schizont is close to full maturity (Fig. 1D). The only dimension on which the extracellular and intracellular merozoites differ is that the extracellular merozoite is spherical while the average intracellular merozoite conforms to a tooth-like shape, likely due to space constraints in the schizont and to maintain a connection to the residual body throughout development. This observation is consistent with data from *Plasmodium knowlesi* and a post PVM-rupture *P. falciparum* schizont^13, 27^. Other characteristics of the ABS are described in comparison with the gametocyte in other sections below.

### Parasite-induced modifications of the red blood cell

RBCs infected with mature gametocytes contain flat, membranous disks (Fig. 2A-B) that differ from the characteristic membranous stacks called Maurer’s clefts we observed in ABS-infected RBCs (Fig. 2C). In gametocytes of both genders, extraparasitic structures were less frequent, generally larger but heterogenous in size and unevenly distributed within the RBC cytoplasm (Fig. 2A-B). We consistently find a higher density of extraparasitic structures in the Laveran’s bib that spans between the two ends of the mature gametocyte^28^. This matches results of previous work that found accumulation of a class of exported proteins to this subcellular location^29^. Furthermore, we find an occurrence of a ‘Garnham body’, a rare gametocyte exclusive structure originally discovered in 1933 with still unknown functional significance^30^ (Fig. S6). Our data confirm the highly membranous appearance of the Garnham body described in previous micrographs and specifically we identify four double membranes. However, we found no associated hemozoin, which was identified in some light microscopy examples^5, 6, 22, 31^ (Fig. S6A-B). The Garnham body was found adjacent to but not within the Laveran’s bib, at a site devoid of IMC and situated in the vicinity of the opening of the cytostome (Fig. S6C). The presence of a Garnham body in the RBC also coincides with electron dense protrusions of the parasite vacuolar membrane radiating in all direction from the parasite and a membranous electron lucent compartment within the parasite, that were observed in no other cells (Fig. S6A-B). These concurrent exclusive features could indicate Garnham bodies to be a sign of an unhealthy gametocyte and/or RBC or suggestive of a rare subtype in gametocyte populations.

Consistent with published literature, the plasma membrane of gametocyte-infected RBCs is not visibly modified, while the ABS-infected cells are thoroughly covered by uniformly distributed knobs, which are proteinaceous parasite-derived protrusions that mediate cytoadherence of ABS-infected RBCs^32^ (Fig. 1C, Movie S3). Reconstructions suggest a total of 390 knobs on the surface of the schizont-infected RBC corresponding to 2.9 knobs/μm^2^, which is in line with previous measurements obtained for NF54-infected RBCs via atomic force microscopy^33^. The overall surface profile of the segmenter-infected RBC appeared more uneven with more peaks and valleys compared to the developing schizont, while gametocyte-infected RBCs appeared to have a smoother surface profile than either (Fig. S5B-D). As anticipated, the trilaminar membrane architecture of the gametocyte-RBC interface, consisting of the parasite vacuolar membrane (PVM), the parasite plasma membrane (PM) and the IMC, is evident from the micrographs. Localized areas of further thickened IMC membrane might represent the leading edge of IMC plates^16^ while areas without IMC cover in immature stages suggest ongoing IMC plate biogenesis (Fig. S7A-B). In mature stages, we occasionally find similar IMC overhangs of unknown significance (Fig. S7C).

### The gametocyte cytostome

RBC-parasite interactions are not restricted to host-cell modification, but also include the internalization and digestion of host-cell cytoplasm to fuel parasite growth. An unusual feeder organelle, the cytostome (meaning cell mouth), facilitates this uptake of RBC cytoplasm in ABS^12^. In our micrographs of ABS and other studies^34^, the cytostome presents itself as a hemoglobin filled tube surrounded by a double-membrane and possessing an electron dense “neck”, called the cytostomal collar, at the invagination site (Fig. S8). Gametocytes similarly rely on hemoglobin internalization and digestion. Putative morphologically diverse cytostomes have been assigned in past EM studies^5, 6^. When trying to assign the cytostome in mature gametocytes, we came across likely yet distinct candidate structures. Presenting a clear invagination of the parasite membranes, the structures are always surrounded by extensive ER but otherwise can be subdivided into three categories (Fig. 3A). In 44% of the evaluated examples (n = 50), the organelle has a membrane delimited electron dense ring that is 100-160nm thick and connected to the RBC cytoplasm while the lumen circumscribed by the ring appears electron lucent and can occasionally contain further membranous or dark stained structures. In 40% of the cases the structures had a more ABS-like appearance with no clear delineation into ring and lumen but circular grooves on the inside of the membrane that are absent in the ABS cytostome potentially indicative of future emergence of the ring structure. In the remainder the ring is relatively larger with inconsistent thickness and a smaller electron lucent internal lumen potentially representing an in-between state. Furthermore, it appears that each gametocyte that we have fully imaged only contains a single cytostome of any of these three gametocyte-specific subtypes. Conversely, in ABS multiple cytostomes have been reported^35^ and are also evident from our data with a maximum of seven cytostomes per cell (Fig. S8, panel 3). Interestingly, in one example of a developing gametocyte, we find both a canonical ABS-type cytostome and a gametocyte variant (Fig. 3B). This observation suggests that the structure we identified in gametocytes might be entirely distinct from the cytostome. In mature stages, hemoglobin internalization is largely finished and consequently the cytostome may no longer be required for its canonical purpose. It is tempting to speculate that this cytostome variant is instead involved in host lipid acquisition, as parasite lipid content dramatically increases throughout gametocytogenesis^36^. This would also offer a plausible explanation for the ER, as the primary site of lipid metabolism, consistently surrounding the cytostome and would suggest that the occasional dark contents of the lumen seen in our preparations are (osmiophilic) lipids.

**Figure 3.**
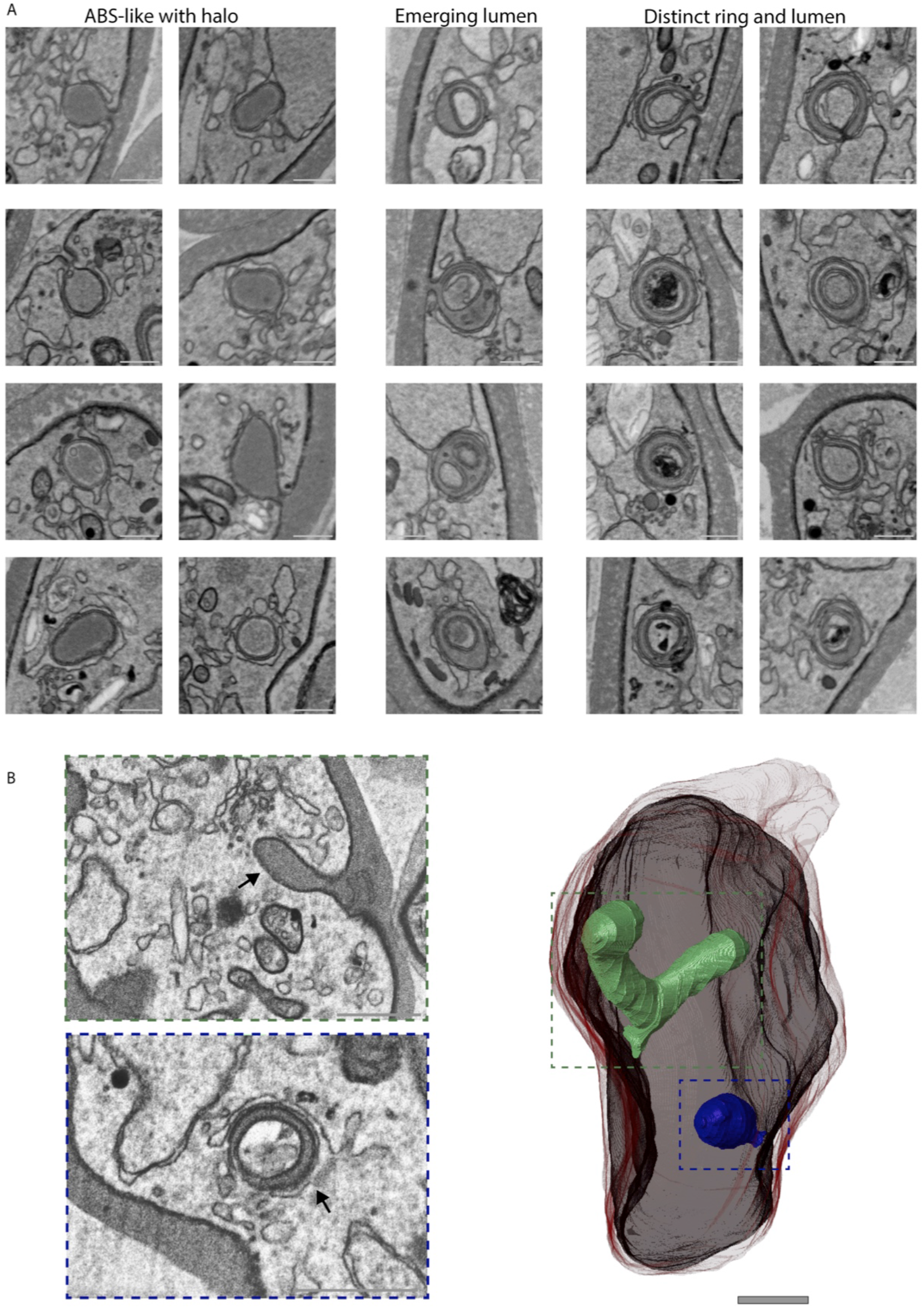
The gametocyte cytostome. **A.** Exemplary micrographs of the cytostome in 20 different gametocytes subdivided into three categories. **Left**: Cytostome without further membrane delineation in lumen, relatively homogenous lumen and circular grooves that could indicate future emergence of ring. **Middle**: Relatively large membrane electron dense ring and small electron lucent lumen. Ring is continuous with RBC cytosol, putative intermediate state. **Right**: Electron dense ring with homogenous thickness and lumen with heterogenous content. Ring is continuous with RBC cytosol. All categories have close association of ER unlike the ABS cytostome. Scale bars = 0.1 μm. **B.** Rendering of a developing gametocyte that contains both a canonical ABS cytostome and a cytostome with distinct ring and lumen. The ABS cytostome (green) is characterized by homogenous dark lumen and continuation of the parasite membrane that forms the invagination. The gametocyte cytostome (blue) is characterized by a bulbous shape and distinct electron dense ring with electron lucent lumen.

### The gametocyte ER and Golgi-apparatus

Universally the ER is an extensive interconnected organelle continuous with the nuclear envelope that serves as a transport hub, hosts various critical cellular functions, is widely connected with other organelles, and even acts as a mediator of organellar interactions^37^. We find that the ER in both developing gametocytes and mature female gametocytes is extensive and occupies large parts of the cell, similar to the schizont (Fig. 1A, Fig. S2, Movie S1). The mature male gametocyte on the other hand appears to contain much less ER, which is in line with previous observations and proteomic data^38, 39^ (Fig. 1B, Movie S2). It is noteworthy that in the mature female gametocyte, the ER is most dense in the polar regions of the parasite. This coincides with the distribution of a closely associated organelle, the Golgi, which, in *Plasmodium*, takes the rudimentary form of dispersed unstacked cisternae and is recognizable as a smooth membraned vesicle cluster^22, 40–42^. In females, the Golgi is predominantly found in two distinct clusters at both polar ends, while in the mature male it appears distributed across smaller clusters, similar to the developing gametocytes that also have a more widespread distribution. Compared to the gametocytes, the Golgi is much more extensive in the schizont and found in evenly dispersed clusters around the cell (Fig. 1C, Movie S3).

### The ER as a putative path across the IMC

The ER also facilitates the extensive host-cell remodeling that occurs in ABS. In *P. falciparum*, an estimated lower bound of 300 unique proteins, >5% of the proteome, is exported to the host cell to modify aspects such as cytoadhesion, the host cytoskeleton, or nutrient permeability^43^. Proteins destined for export are processed in the ER and delivered to the PM via vesicular transport and then forwarded to the *Plasmodium* Translocon of EXported proteins (PTEX) in the PVM, which translocates the proteins into the host cell. While extensively researched in ABS, we know relatively little about how this process translates to the gametocyte situation. Maurer’s clefts appear absent in gametocytes, possibly replaced by other cleft-like structures^44^, and the gametocyte exportome is distinct from the ABS exportome^45^. Whilst protein export machinery is essential for early gametocyte development^46, 47^, protein export has not been conclusively demonstrated in gametocytes beyond stage III. Indeed, as the IMC cover increases, staining of the export machinery on the PVM becomes less pronounced or even disappears completely in *P. falciparum* and *P. berghei* gametocytes, respectively^45, 48^. If protein export still happens in late gametocyte stages the mechanism by which the additional barrier presented by the IMC is overcome, is currently unknown^49^. In mature gametocytes, we frequently find sites at which the ER runs very close to or is in direct contact with the parasite membrane (Fig. S9). From our observations this interaction can manifest in a few different forms. The most frequent observation is that ER directly contacts the IMC, leading to a continuum between these two compartments as well as local disruption of the IMC (Fig. S9A). Less frequently but consistently, we observe “budding” of a piece of ER that contacts the parasite membrane (Fig. S9B). These local continuities between ER and IMC might represent a mechanism to bridge the IMC and allow canonical ER-derived vesicle fusion with the PM or facilitate alternative means of protein export. In developing gametocytes with incomplete IMC cover, we find that all extension sites are subject of extensive ER interaction with the nascent IMC (Fig. S9D) This is in line with previous EM-based observations that the outer nuclear envelope interacts with nascent IMC and the PPM in developing gametocytes^15^ and with immunofluorescence microscopy data suggesting regions of ER contact with the IMC^50^. Even in mature gametocytes with complete IMC cover we too find sites at which ER converts to electron dense material that resembles IMC (Fig. S9C). Taken together, these observations, while anecdotal, provide further evidence for a dynamic interplay between ER and IMC biogenesis or maintenance and suggest a speculative mechanism to bridge the additional membrane layer in gametocytes for protein export.

### Features of *Plasmodium* mitochondria

In gametocytes, the mitochondria are readily recognizable based on their cristate appearance and two-layered membrane. The cristae appear to have some degree of interconnectivity, consistent with previous interpretations of tubular cristae, but the small size of the structures and variability between slices, even at a z-resolution of 15 nm, makes clear assertions of connectivity challenging (Fig. S10B, Movies S4-5). While edges of the individual cristae are more evident from the lower noise serial sectioning data, the low z-resolution does not allow for reconstruction of their morphology (Fig. S10A). In the individual micrographs, the cristae are similar to the recently described bulbous cristae in the related apicomplexan parasite *Toxoplasma gondii*^51^. In line with previous reports, the mitochondrion is double-membraned yet acristate in ABS. In both ABS and gametocyte mitochondria, we regularly identify electron dense mitochondrial granules (EDMGs) that appear in clusters of 1-10 EDMGs and are heterogeneous in volume with a notably smaller size, propensity to be more separate and more slender in shape in ABS (Fig. 4A) ^52, 53^. While they have not been described before, they are readily recognizable in the image stacks of schizonts made public by Rudlaff *et al*.^13^, indicating it is not a unique feature of our preparations. The electron dense appearance could suggest that these granules are sub-compartments that are either very proteinaceous or contain polysaccharides, lipids, or mitochondrially relevant metals such zinc, calcium, copper, and/or iron. *P. falciparum* mitochondria have been shown to play a role in calcium mobilization and storage^54^ and so-called matrix granules that are heavily enriched in calcium phosphate have been found in mammalian mitochondria^55^. In parasites isolated from fish digestive tracts, similar mitochondrial granules have been shown to store glycogen, which our staining procedure is particularly well suited to show^56, 57^. Increase of energy storing granules in gametocyte stages could be a plausible preadaptation for survival in the relatively nutrient deprived environment of the mosquito midgut. Alternatively, the EDMGs could represent the mitochondrial RNA granules described in mammalian mitochondria. These are highly proteinaceous, non-membrane delimited sub-compartments in the mitochondrial matrix that are thought to play an important role in mitochondrial RNA processing, mitoribosome assembly, and gene expression^58^. As mitochondrial translation in *Plasmodium* purely relates to complex III and complex IV components of the respiratory chain, this could explain the discrepancy in EDMG size between the comparatively respiratory chain-poor ABS compared to the respiratory chain-rich gametocytes^20, 59^.

**Figure 4.**
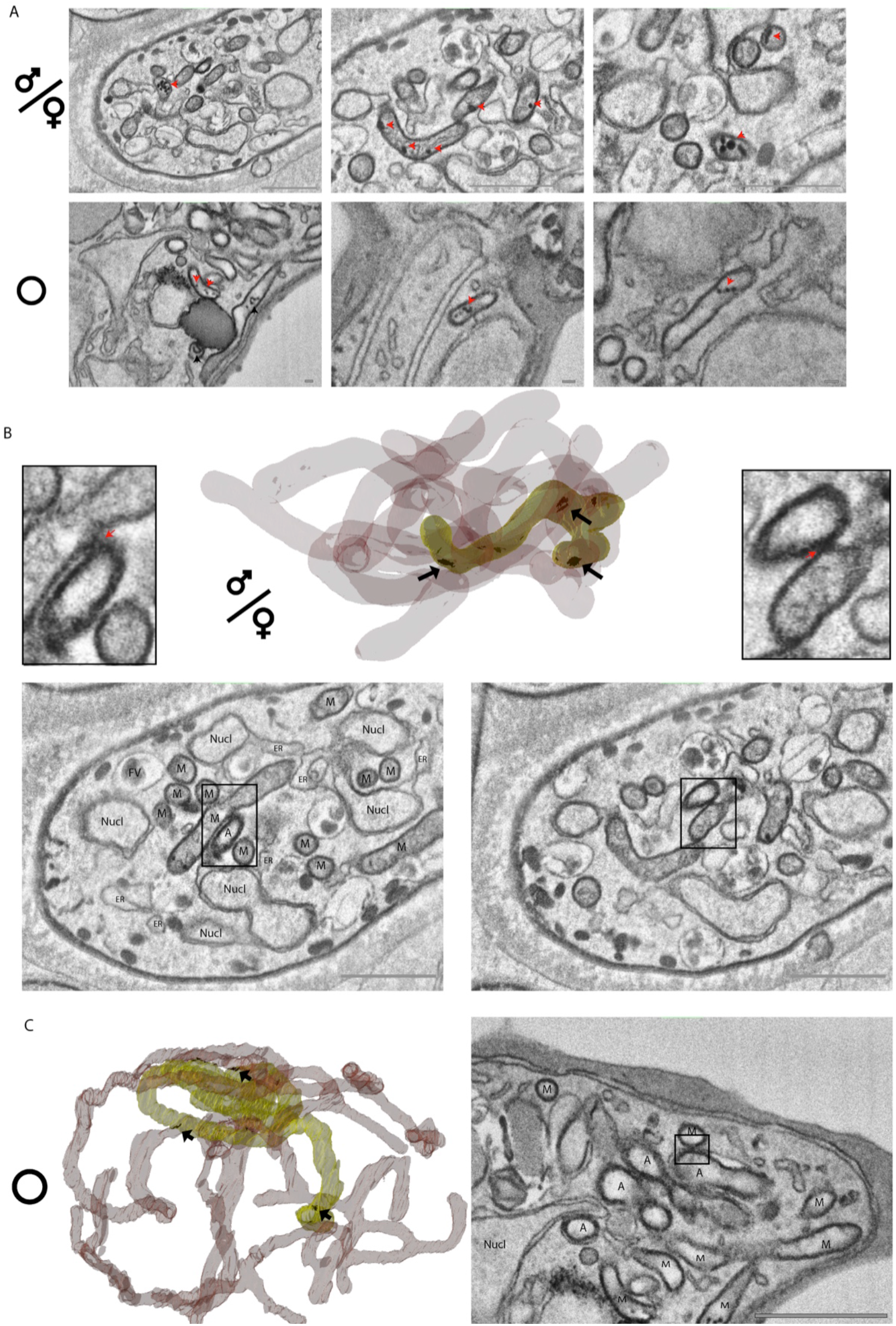
Mitochondrial vesicles and interaction with the apicoplast. **A.** Representative micrographs of EDMVs in gametocytes (upper panel, scale bars = 1 μm) and asexual blood stages (lower panel, scale bars = 0.1 μm). Arrowheads highlight examples of EDMVs. **B.** Rendering of exemplary gametocyte mitochondrion (red, high transparency) and apicoplast (yellow, low transparency) with putative organelle interfaces rendered in black. Representative micrographs for two interfaces and respective zoomed crops are shown below. Scale bars = 1 μm. **C.** Same representation as panel B for a schizont. M = Mitochondrion; ER = Endoplasmic reticulum; Nucl = Nucleus; FV = Food vacuole.

### Multiple mitochondria in disguise

The current consensus is that, like ABS, gametocytes contain a single heavily branched mitochondrion with diverse morphology that does not closely correspond to gametocyte sex or maturity^19^. Hence, we were surprised to find that while branched and clustered, we did not encounter a fully interconnected mitochondrion in our imaged gametocytes. Instead, we found multiple mitochondria (5-10) per cell that, while clustered and with close membrane appositions, did not share a continuous lumen (Fig. 5A-B, Movies S5-6). To investigate this further, we carefully revisited mitochondrial morphology of a larger number of mature gametocytes using fluorescence microscopy. As expected, most images gave the impression of a fully interconnected organelle, but we also found examples where at least some mitochondrial staining appeared separate from the major cluster (Fig. 5C, Movies S7-10). This is consistent with the FIB-SEM data, where completely separated mitochondria can be observed (Fig. S11A), though in most cases the mitochondria are in close proximity, potentially even suggestive of homotypic membrane interactions that are commonplace among fungal and mammalian mitochondria^60^ (Fig. 5B). These short distances of <50nm are well below the diffraction limit of conventional fluorescence microscopy and would appear as a continuous organelle. We observed no morphological indications for poor gametocyte health, which was confirmed by normal gamete activation and the ability of male exflagellation. Moreover, exflagellating males have clearly distinct, dispersed, and rounded mitochondria (Fig. 5D, Movies S11-12). In our FIB-SEM data, we identified one comparable example of a male gametocyte showing bloated, rounded, and dispersed mitochondria that appeared to be in the process of separating, though at this point it is unclear whether this is a male gametocyte at the onset of activation or an aberrant cell (Fig. S11B). We find occasional mature male gametocytes showing a similar pattern using fluorescence microscopy (Fig. S11C). Generally, cristae appeared homogenous across the multiple mitochondria within a gametocyte and were also observed in the bloated and dispersed phenotype (Fig. S11B,D). In comparison, the asexual schizont possesses a single mitochondrion with a continuous lumen, consistent with previous observations, suggesting that the observed phenotype is not directly caused by fixation or sample processing (Fig. 1C) ^13^. Taken together, we find consistent evidence for deviation from the current consensus of a fully interconnected mitochondrion in gametocytes. The functional significance of this deviation and appearance in other strains and culturing systems remain to be investigated.

**Figure 5.**
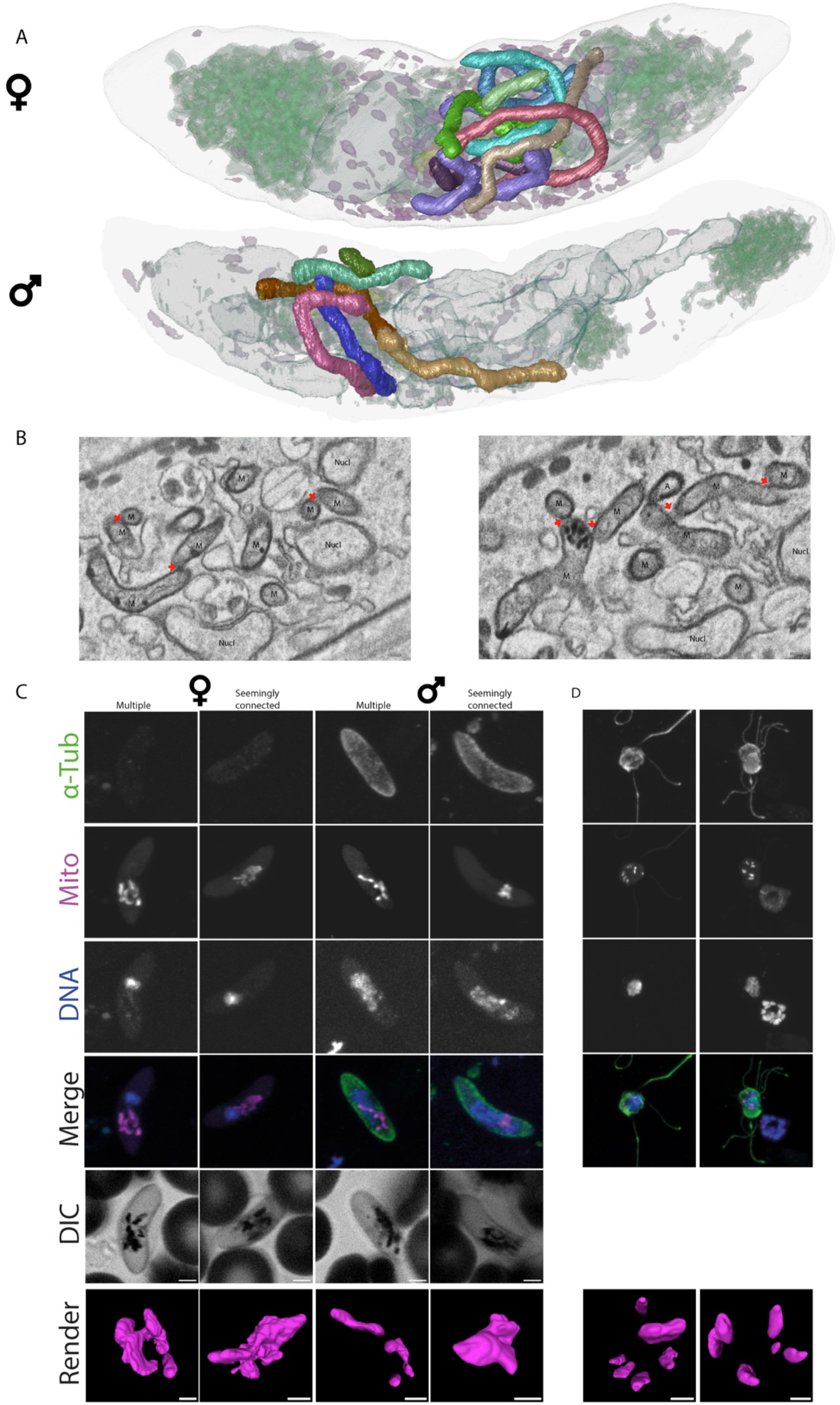
Gametocytes contain multiple mitochondria in close vicinity. **A.** Rendering of mature female (upper panel) and male gametocyte (lower panel). Each mitochondrion is rendered in a separate color. The male gametocyte has six distinct mitochondria while the female gametocyte has nine distinct mitochondria. In the female gametocyte membrane appositions are more frequent and mitochondria are more closely associated. Nucleus, osmiophilic bodies, ER, Golgi, and apicoplast are rendered with high transparency to provide cellular context. **B**. Exemplary micrographs showcasing membrane apposition between different mitochondria indicated by red arrows. The mitochondrial membrane remains intact at these sites and no continuity between the different mitochondria is evident. Scale bar = 0.1 μm. M = Mitochondrion; A = Apicoplast; Nucl = Nucleus. **C.** Immunofluorescence analysis of mature and female gametocytes. Depicted left to right are maximum intensity projections using anti-α-tubulin antibodies, MitoTracker^™^, DAPI, merge of all channels and differential interference contrast (DIC) images. α-tubulin was used to distinguish male and female gametocytes, DNA was visualized using DAPI and mitochondria were visualized using MitoTracker^™^. For each gender one example was given for recognizably separate mitochondria and a seemingly interconnected mitochondrion. Scale bar = 2 μm Additional renderings of the mitochondrion were generated based on the mitochondrial fluorescence signal (bottom row, scalebar = 1 μm). **D.** Immunofluorescence analysis of exflagellating male gametes. Channels are the same as in **C.** Mitochondria are clearly separate and dispersed throughout the cell body.

### Appositions of mitochondrion and apicoplast

In gametocytes, the mitochondria form a network that is wrapped around the apicoplast (Fig. 1 A-B, Movies S1-2). The apicoplast is recognizable as a clearly distinct structure from the mitochondrion due to the thicker appearance of its four surrounding membranes and lack of internal membranous structures and more electron lucent lumen. The mitochondrial-apicoplast network is not spread throughout the whole cell but localized to a relatively central area of the gametocyte, always adjacent to the nucleus. It is noteworthy that this association and the mitochondrial network are much tighter than what we observe in schizonts where the mitochondrion permeates the whole cell (Fig. 1C, Movie S3). The tight association is clearly distinct from the spatial separation of these organelles in sporozoites but akin to the tight association that has been observed in liver-stage parasites^61, 62^. Next to a general vicinity of the two organelles, we observe electron dense junctions spanning the membranes of the two organelles in both gametocyte and ABS (Fig. 4B-C). While it remains unresolved whether these sites constitute a true MCS (Information Box 1), the thickened electron dense interaction area is indicative of a tethering structure, providing a reasonable explanation why the organelles are consistently co-purified^63^. While a putative physical connection does not provide mechanistic evidence, the observed phenomenon may facilitate metabolic cooperation, *e.g*. by serving as an exchange site for metabolites of the heme biosynthesis pathway for which the enzymes are localized partly in the mitochondrion and partly in the apicoplast^64, 65^.

#### Textbox 1 Membrane contact sites

Membrane contact sites (MCS) have typically been defined as areas of membranes of two organelles that are in close proximity to each other. These areas can be homotypic (between the same type of organelle, e.g. mitochondrion-mitochondrion MCS) or heterotypic (between different organelle types, e.g. mitochondrion-ER MCS). To increase clarity and create a unified vocabulary Scorrano *et al*.^87^ have put together a set of unifying features that a MCS should have to be characterized as such.

1. There should be tethering forces from either protein-lipid or protein-protein interactions that maintain the distance. This replaces previous proximity ranges that were used as guidelines as contact sites have been found that far exceed these distance guidelines.
2. There should be no fusion of membranes. If there is fusion of membranes this can be referred to as docking or fusion. Vesicular transport at the contact site may exist within this definition.
3. All contact sites should fulfill a specific function. A disruption of a contact site should impact cell function.
4. MCS should have a defined proteome or lipidome that is required for their function, maintenance, or regulation.
5. Time is not a factor. Dynamic or transient structures that fulfill the other requirements can be considered a MCS.

### Appositions of ER and mitochondrion

From mammalian and yeast mitochondria, we know that the ER is intimately linked to and interacts with the mitochondrion through MCS. In these species, MCS cover 2-5% of the mitochondrial surface area and are composed of a known set of interactive proteins^66, 67^. These sites are thought to be crucial for lipid homeostasis and calcium transport. In our data, we similarly find multiple sites where the ER is in very close proximity to the mitochondrion (Fig. 6C-D). Anecdotally, both in the schizonts and gametocytes, we find examples where these sites are accompanied by EDMGs appearing to span the mitochondrial membrane and contact the ER (Fig. S12), which provides tentative support for a role of EDMGs in calcium storage and/or mobilization. The appearance of the ER-mitochondrion appositions resembles micrographs of previously described ER-mitochondrion MCS^68^, leading us to believe that they are conserved in *Plasmodium*. However, from all previously described tethering complexes, the *Plasmodium* genome is lacking at least one critical component^69–72^. In *S. cerevisiae*, the ER membrane complex (EMC) and TOM5 facilitate phospholipid exchange between ER and mitochondrion^73^. Recently, we demonstrated that an assembled EMC is present in *P. falciparum*^20^, but a TOM5 orthologue is not encoded in *Plasmodium*. This prompted us to re-interrogate the underlying complexomics data and look for another potential mitochondrial interactor with the EMC. In doing so, we found that *Pf*TOM7, another component of the translocase of the outer membrane (TOM) complex, comigrates in native electrophoresis with the EMC as well as the TOM complex (Fig. S13A). Looking at the multiple sequence alignment, few residues are conserved outside of the transmembrane region and none are shared between *Sc*TOM5 and *Pf*TOM7 (Fig. S13B-C). While overall structural features of *PfTOM7* and *Sc*TOM7, based on alignment of the *Pf*TOM7 AlphaFold prediction^74^ and experimentally determined structure of *Sc*TOM7^75, 76^ appear similar, *Sc*TOM5 is expectedly less similar outside of the (predicted) TM helix (Fig. S13D). This suggests that if *Pf*TOM7, based on comigration with the EMC complex, is a tether, the specific binding site is unlikely to be conserved. If a different mechanism tethers these two complexes, it might also suggest that there is biological utility in specifically facilitating the tether through a component of the TOM complex.

**Figure 6.**
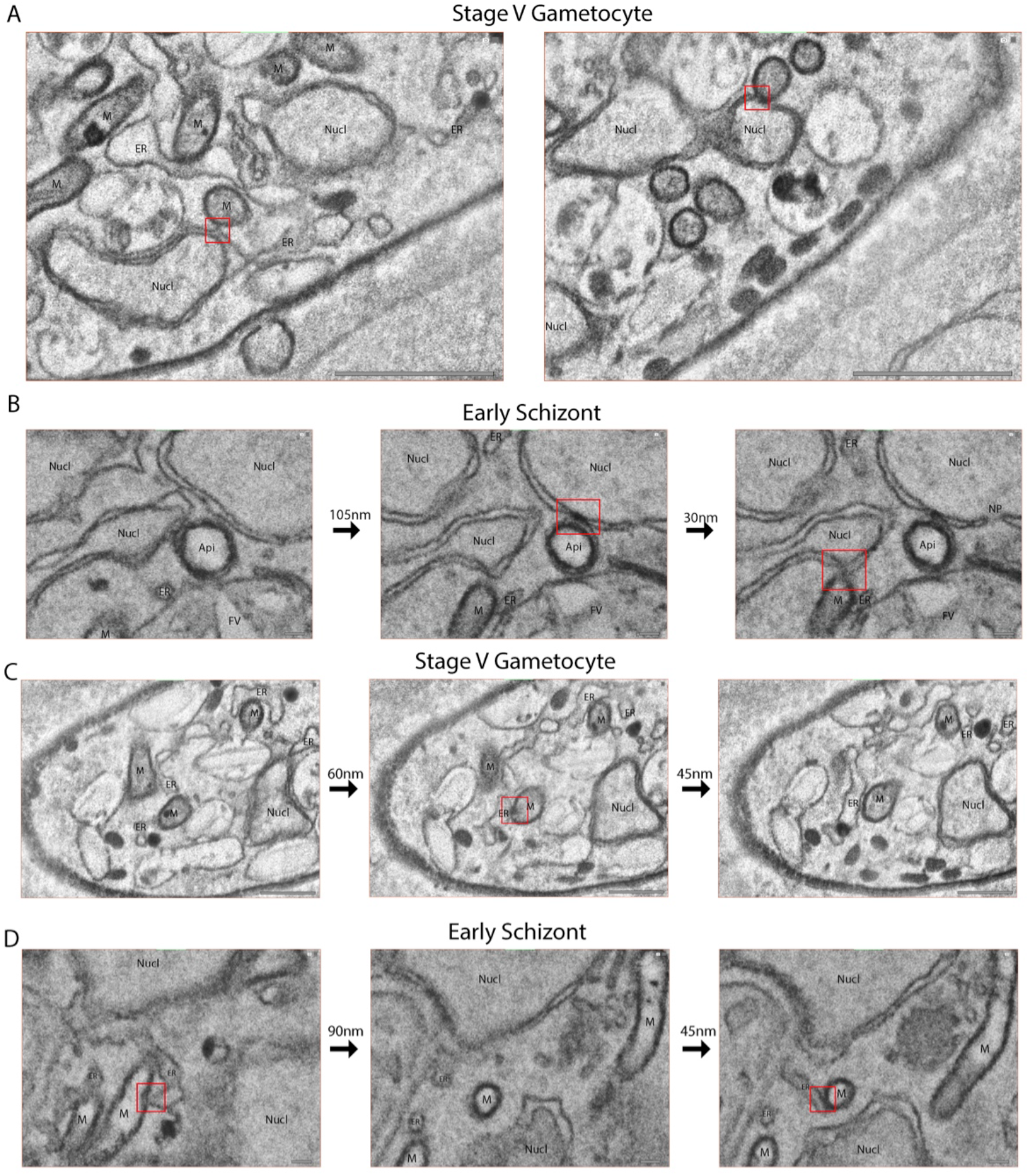
Putative organelle interfaces in blood stages. **A.** Representative micrographs showing putative mitochondrion-nucleus interaction sites in a mature gametocyte. The respective sites are highlighted through a red box. Scale bars = 1 μm. **B.** Representative series of micrographs showing putative mitochondrion-nucleus and apicoplast-nucleus interaction sites in a schizont. The three frames are crops at the same x,y-coordinates in the acquisition plane but are at different z-depth with the respective distance displayed between micrographs. **C.** As in (B), showing putative mitochondrion-ER interaction sites in a mature gametocyte. Scale bars = 1 μm. **D.** As in (B), showing putative mitochondrion-ER interaction sites in a schizont. Scale bars = 0.1 μm M = Mitochondrion; ER = Endoplasmic reticulum; Nucl = Nucleus; FV = Food vacuole.

### Features and dimorphism of the gametocyte nucleus

The nucleus is bounded by an inner and outer leaflet of the nuclear envelope, where the outer leaflet is continuous with the ER and the inner leaflet circumscribes the nucleus. In gametocytes, we find an intriguing deviation from the typically round or oval nuclear shape^77^. The nuclei share a bulbous body from which thinner diverse extensions extrude predominantly in one direction covering large parts of the cell (Fig. 7A-B). The bulbous body always contains a more electron dense, not membrane-bound region, which has previously been assigned as the nucleolus of the female gametocyte^22^ (Fig. 7C). The observed nuclear shapes are largely consistent with prior studies that noted a discrepancy between a relatively small and localized DNA stain and an elongated staining from a nucleus-targeted fluorophore^15, 78^. In stage IV – V gametocytes, a clear nuclear dimorphism is apparent with female gametocytes possessing a less complex smaller nucleus with an average volume of 6.6 ± 0.3 μm^3^, while males have a more complex nuclear shape and a much higher nuclear volume at 12.4 ± 1.1 μm^3^ (Fig. 7B). A comparatively enlarged nucleus in males is consistent with well-known differences observed in Giemsa-stained samples and might prepare male gametocytes for the rapid nuclear division and DNA replication that they undergo upon activation. While smaller, the female nucleus is still relatively large, possibly explained by the proposed role of the nuclear extension in elongation of the gametocyte^15^. Based on our measurements, for merozoites the same genetic material fits into a nucleus of ~0.5 μm^3^ while for our exemplary schizont in the process of DNA duplication, average nuclear volume is around 1.8 μm^3^. Stage II - III immature gametocytes also reflect the nuclear dimorphism observed in stage IV – V gametocytes, with some having a relatively smaller and less complex nucleus while another subset has a bigger and more complex nucleus (Fig. 7A). While we cannot confidently assign the developing gametocytes to either gender, a plausible hypothesis would be that the subset with smaller nucleus represents immature female gametocytes, while cells with larger nucleus represent immature male gametocytes. However, this is in conflict with a recent finding that commitment, as indicated by measurable differences in gene expression, to either male or female cell fate is only decided at stage III^79^. Furthermore, a recent study utilizing array tomography has found no such nuclear dimorphism in earlier stages but does show the same similar dimorphism and absolute volumes to our findings for later stages^15^. It would be interesting to see whether these different observations stem from differences in experimental methods, stage assignment, image segmentation or biological material.

**Figure 7.**
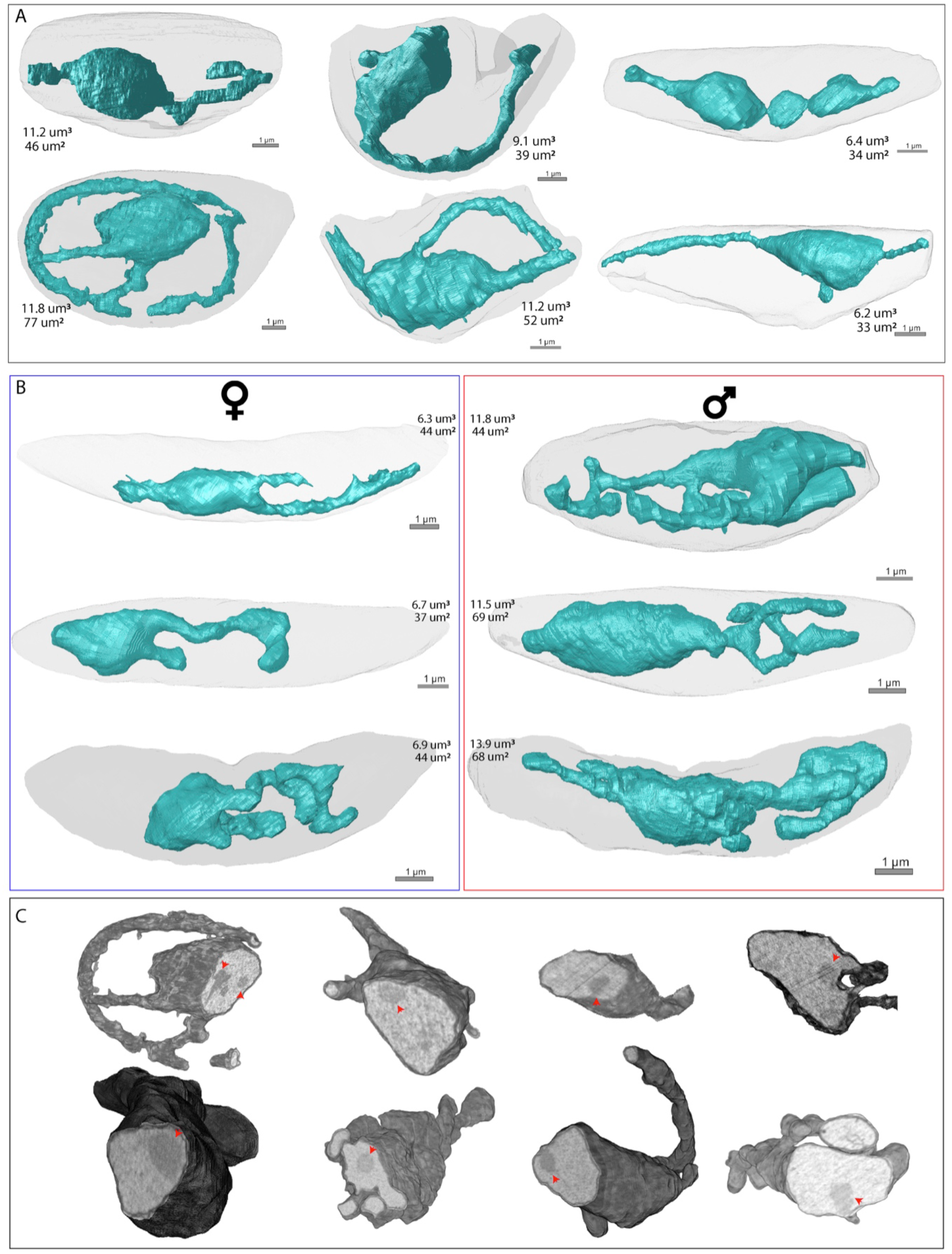
Distinct nuclear morphology in gametocytes. Renderings of nuclei (teal) in (**A**) stage II-III and (**B**) stage IV-V gametocytes. Stage IV/V gametocytes are divided in female (blue outline) and male (red outline) gametocytes based on the number and appearance of osmiophilic bodies, hemozoin distribution and ER prevalence. For all nuclei, the respective volumes and surface areas are indicated. **C.** Rendering of cross section of nuclei with gray values from EM data overlaid. Red arrowheads point at the position of the putative nucleolus.

### Appositions of nucleus and endosymbiotic organelles

In *Plasmodium*, both mitochondrion and apicoplast have essential organellar ribosomes, with promising features as drug targets^80–83^. Nevertheless, neither have an associated set of organelle targeted tRNAs that would be sufficient to facilitate translation of their respective genomes^84^. Recently, it has been found that the nuclear-mitochondrial MCS play a role in RNA exchange and signaling. These sites are distinct from the well-characterized peripheral ER-mitochondrial MCS but likewise enabled by tether proteins without obvious homologues in the *Plasmodium* genome^85, 86^. We find distinct appositions of the nucleus and mitochondrion in both schizonts and gametocytes (Fig. 6A-B). At these sites there appears to be continuity of the outer leaflet of the nuclear envelope and the outer mitochondrial membrane. In contrast, we find that the nucleus-apicoplast interface in ABS is characterized by a local condensation of the normally spatially separated layers of the nuclear envelope into one electron denser layer without obvious membrane fusion (Fig. 6B). Whereas the mitochondrion in gametocytes closely envelops large areas of the apicoplast, only a subset of cells have close physical proximity between nucleus and apicoplast and in those cells we never find indications of direct membrane contact. Functionally, interfaces of apicoplast and mitochondrion with the nucleus might serve phospholipid homeostasis or RNA exchange as has been suggested in model eukaryotes^85, 86^, or provide a potential mechanism for import of organellar tRNAs without known targeting signals thus shoring up the incomplete translational machinery in these organelles.

## Conclusion & Future Outlook

In this study, we applied volumetric electron microscopy to the transmittable gametocyte stages of the malaria parasite, allowing us to place findings from the past into their current molecular context and tackle questions that are challenging to address using single sections or serial sections with low z-resolution such as the connectivity within or between various organelles. Furthermore, these data allowed us to create the first 3D visualization of Golgi, gametocyte cytostome, Garnham body, and ER in gametocytes, improving our understanding of the cellular architecture in this part of the life cycle. In doing so, we are aware of the descriptive nature of this study and the limitations of the tool applied. Any of the hypotheses put forward are meant to provide a foundation for controlled molecular studies and targeted approaches that are more suitable to identify the underlying molecular players.

### Public access and potential for reusability

FIB-SEM is a powerful tool, particularly for a cellular system that is as unusual as malaria parasites, as it allows determination of general ultrastructural organization and basic measurements to cover knowledge gaps that in other model eukaryotes have long been closed. Similar to other global datasets, one of the biggest challenges is to extract all relevant information and leverage the utility of non-targeted approaches and not just use them to answer one narrow question. To facilitate future re-interrogation of the data, we have deposited all underlying image stacks, as well as all analyzed cells and their corresponding segmentations, in the Electron Microscopy Public Image Archive (EMPIAR-11497). We hope that the latter can serve as training data for machine learning by groups with more suitable hardware/software and deep learning expertise to greatly simplify annotation workflows in the future.

## Materials & Methods

### Parasite culture & gametocyte induction

All parasite material used in this study is derived from the NF54/iGP2 strain. NF54/iGP2 parasites were previously shown to be able successfully complete the whole lifecycle as well as not markedly differ from wildtype NF54 parasites in other parameters that were investigated^78^. This parasite strain has the desirable property that gametocytogenesis can be selectively induced by removal of glucosamine from the medium, which triggers sexual commitment through the transcriptional cascade centered around the AP2G gene^88^. Asexual blood-stage parasites (ABS) were maintained according to standard culturing procedure in RPMI [7.4] supplemented with 10% human serum and 5% hematocrit using standard culturing technique in a semi-automatic culturing system^5, 89^. For the maintenance of the NF54/iGP2 strain, an additional supplement of 2.5 mM D-(+)-glucosamine hydrochloride (Sigma #1514) was used. To induce gametocytogenesis, glucosamine was omitted from the culturing medium. Between days 4 and 8 after gametocyte induction 50 mM N-acetylglucosamine (Sigma #A6525) was used to eliminate ABS parasites. On day 14 gametocyte-infected RBCs were separated from uninfected RBCs through magnetic separation as described previously^90^. Close attention was paid to prewarm and maintain all solutions and apparatus at 37°C to avoid unintended activation of mature gametocytes. As reference material trophozoite and schizont stages from mixed ABS cultures were similarly enriched through magnetic separation and processed alongside the gametocyte cultures. For sample 3, N-acetylglucosamine treatment was omitted to control for phenotypes causes by this treatment. This omission leads to presence of ABS in the resulting images (sample 3).

### Sample preparation for electron microscopy

Samples were prepared as described previously^20^. Briefly, the enriched infected red blood cells were fixed using 2% glutaraldehyde in 0.1 M cacodylate (pH 7.4) buffer overnight at a temperature of 4 °C. The fixed cells were then washed and the cell-pellet was resuspended in 3% ultra-low-gelling agarose, solidified, and cut into small blocks. The agarose blocks containing the fixed cells were postfixed for 1 hour at room temperature using a solution of 2% osmium tetroxide and 1.5% potassium ferrocyanide in 0.1 M cacodylate buffer containing 2 mM CaCl2, washed in MQ and then treated with 0.5% thiocarbohydrazide for 30 minutes at room temperature. After washing, the agarose blocks were again suspended in 2% osmium for 30 minutes at room temperature, washed, and then placed in a 2% aqueous uranyl acetate solution overnight at 4 °C. The blocks were then washed and placed in a lead aspartate solution (pH 5.5) for 30 minutes at 60 °C, washed, dehydrated using an ascending series of aqueous ethanol solutions, and subsequently transferred via a mixture of acetone and Durcupan to pure Durcupan (Sigma) as an embedding medium. The aforementioned staining procedure is primarily optimized for high membrane contrast and staining of lipid-rich structures due by utilizing the combination of potassium ferrocyanide, osmium tetroxide and thiocarbohydrazide^91^. Uranyl acetate and lead aspartate furthermore stain DNA/RNA, proteins as well as carbohydrates^92, 93^.

### FIB-SEM

After polymerization and in order to create a smooth plane for FIB-SEM imaging, the sample surface was smoothened using an ultra-microtome (Reichert Ultracut S ultramicrotome (Leica microsystems)). The region of interest was cut out of the resin block and glued on an SEM stub with carbon tape, using conductive silver paint. The samples were then coated with a gold sputter coater (Edwards, Stockholm, Sweden) before introduction into a Zeiss Crossbeam 550 FIB-SEM (Carl Zeiss).

Multiple coarse trenches were milled using a 30 kV@30 nA probe to choose the regions of interest for further 3D volume imaging. Parameters for serial sectioning imaging of large regions were set using the Atlas 3D software (Atlas Engine v5.3.3). The large trenches were first smoothened using a 30 kV@1.5 nA FIB probe, and thereafter a 30 kV@700 pA probe current was used for serial FIB milling. InLens secondary and backscattered electron microscopy images were simultaneously collected at an acceleration voltage of 2.0 kV with a probe current of 500 pA. The backscattered grid was set to −902 V. For noise reduction, images were acquired using line average (n=1) and a dwell time of 3,0 μs. The milling and imaging processes were continuously repeated and long series of images were acquired.

**Table 1.** Basic parameters of FIB-SEM stacks used in these studies. All samples constitute biological replicates.

|  | Sample 1 | Sample 2 | Sample 3 | Sample 4 | Sample 5 |
| --- | --- | --- | --- | --- | --- |
| Life-cycle stage | GCT | GCT | GCT w/ ABS | ABS | ABS |
| Dimensions (x,y,z) [nm] | 4268x2784x2293 | 4428x3804x952 | 4402x2850x900 | 4423x1430 x 1085 | 4010x3157x739 |
| Total Volume Imaged [μm<sup>3</sup>] | 10217 | 6013 | 4234 | 2573 | 3508 |

### Serial sectioning

To have a lower noise reference and identify potential FIB imaging artifacts one gametocyte sample was also analyzed through serial sectioning combined with SEM. After checking for desired density of cells with toluidine blue corresponding resin block was trimmed to the desired size and mounted onto the chuck of a Leica Artos 3D Ultramicrotome. The block was sectioned into thin 80nm slices and sections were collected onto Indium Tin Oxide glass coated with 5 nm carbon. The sections were then imaged with a scanning electron microscope (Sigma300, Zeiss) at an acceleration voltage of 30 kV (HDBSD, 60 μm, high current) using Atlas 5 software. In total 47 consecutive images were taken leading to a total imaged volume of 15124 um^3^.

### Post-processing and segmentation

All processing, visualization and analysis performed in the ORS Dragonfly software (V2022.2). Wherever necessary image stacks were aligned using the mutual information and sum of squared differences registration method and results were manually controlled and adjusted whenever necessary. Contrast was enhanced through application of contrast limited adaptive histogram equalization (CLAHE). 3D segmentation was performed using either manual segmentation or deep learning based segmentation, based on which cellular feature was segmented. Deep-learning based segmentations were controlled for errors and manually adjusted when necessary. For 3D rendering the segmented regions of interest were converted to triangle meshes.

### Immunofluorescence assays

To induce gametocytes, NF54 parasites were swapped from their earlier described complete culture media to RPMI media supplemented with 0.5% Albumax (AlbuMAX II^™^, Thermo Fisher, #11021-037) at 26 hours post invasion. The parasites were allowed to commit for 36 hours, before being put back on complete culture media. At day 12 post induction, stage V gametocytes were stained in 100 nM MitoTracker^™^ (MitoTracker^™^ Orange CMTMRos, Thermo Fisher, #M7510) diluted in complete media at 37°C for 30 minutes. Stage V gametocyte samples were diluted 1:10 in warm complete media before being allowed to settle on pre-heated Poly-L-Lysine coated coverslips (Corning, #354085) for 15 minutes at 37°C, while activated gamete samples were mixed 1:1 with in 30μM xanthurenic acid (Sigma Aldrich, #D120804) and subsequently settled for 15 minutes at RT. Both samples were fixed with 4% paraformaldehyde (Thermo Fisher, #28906)) and 0.0075% glutaraldehyde (Panreac, #A0589,0010) in PBS for 20 minutes at RT. The fixed samples were, permeabilised with 0.1% Triton X-100 in PBS for 10 minutes and blocked in 3% BSA (Sigma Aldrich, #A9418) for 1 hour at RT. To differentiate between male and female gametocytes, the samples were stained with primary α-Tubulin antibody (Thermo Fisher, #MA1-19162) for 1 hour at RT, which was visualised with Donkey anti-Mouse Alexa Fluor^™^ 647 (Thermo Fisher, #A-31571). Finally, samples were stained with 300 nM DAPI (Thermo Fisher, #62248) for 30 minutes at RT before being mounted on a microscope slide using VECTASHIELD (VWR, #H-1000) and sealed in nail polish. The samples were imaged on an confocal LSM900 microscope with airyscan (Zeiss). Z-stack images were obtained with a 0.14 μm step size. Laser power and detector sensibility are maintained the same throughout all the images. A maximum projection was created for every image, in which the tubulin signal was pre-set in order to differentiate between male and female gametocytes.

## Acknowledgements

We thank the molecular team members of the Malaria Research Group for fruitful discussions. We also thank Nico Sommerdijk and the Radboud Technology Center Microscopy - Electron Microscopy Center for supporting this work and Sabrina Absalon for critically proofreading the manuscript. F.E. and T.W.A.K. were supported by the Netherlands Organisation for Scientific Research (NWO-VIDI 864.13.009), C.B. by a PhD fellowship from the Radboud Institute for Molecular Life Sciences, Radboudumc (RIMLS018-009b), J.M.J.V. was supported by an individual Radboudumc Master-PhD grant. R.R. and A.A. are supported by an ERC Advanced Investigator grant (H2020-ERC-2017-ADV-788982-COLMIN for Nico Sommerdijk). A.A. is also supported by the NWO (VI.Veni.192.094).

## Author contributions

F.E., R.R, C.B., M.K.L and J.M.J.V. performed experiments. R.E.S provided conceptual advice and analysis. F.E. analyzed the results, prepared illustrations and wrote the manuscript draft. A.A. provided resources technical expertise and conceptual advice. T.W.A.K. contributed to conceptual development, provided resources and edited the manuscript. All authors contributed to data interpretation and provided feedback on the manuscript. All authors approved the final version of the manuscript.

## Notes

### Competing Interest Statement

The authors have declared no competing interest.

### Summary of Updates

One of our most surprising findings refutes the long standing dogma that malaria parasites have only a single mitochondrion throughout their life cycle. In order to further substantiate our finding that Plasmodium falciparum sexual blood stages have multiple tightly linked mitochondria, we revisited fluorescence microscopy analysis of these stages and included it in the paper.

