## Supplementary Information for "Comparative 3D ultrastructure of *Plasmodium falciparum* gametocytes"

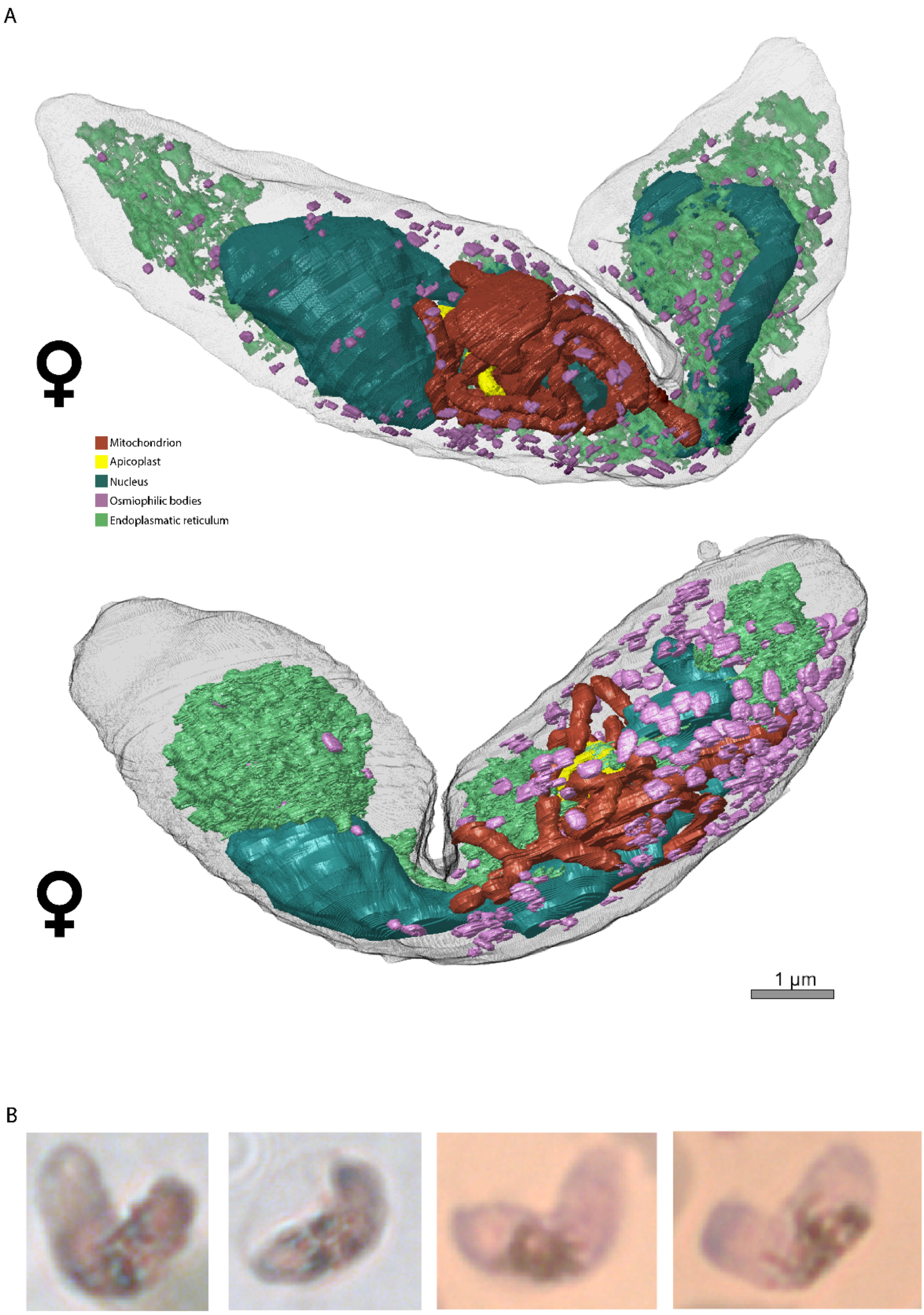


Figure S1. **Ultrastructural features of contorted mature female gametocytes.**
**A.** Renderings of two contorted mature female gametocytes with a typical contorted appearance observed in gametocyte cultures. Ultrastructural features shown include nucleus, mitochondrion, ER, Golgi, osmiophilic bodies, and apicoplast. **B.** Exemplary crops from Giemsa-stained smears of gametocyte cultures showing mature female gametocytes with similar knicked appearances.

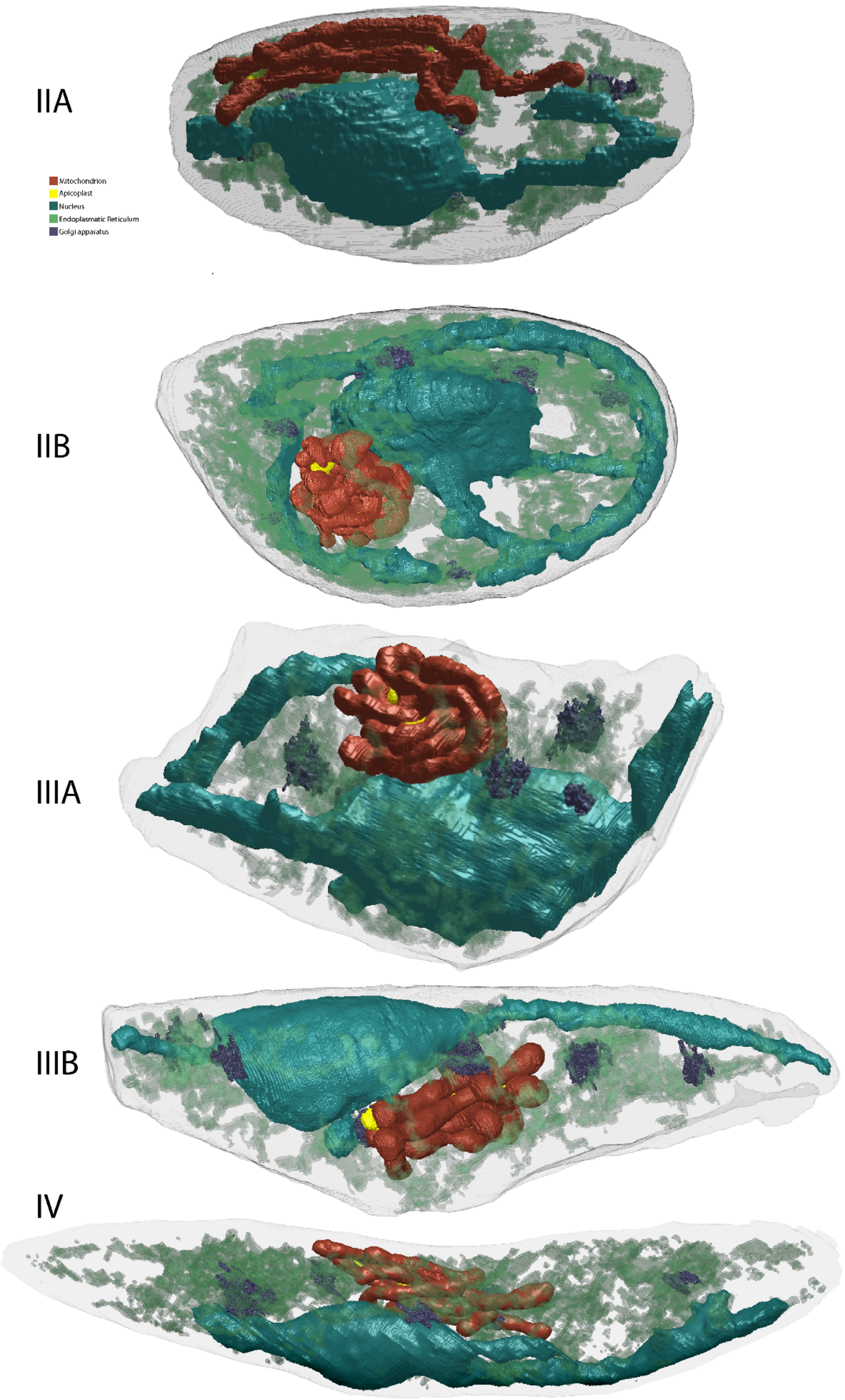


Figure S2. **Ultrastructural features of gametocytes in different developmental stages.**
Renderings of gametocytes in different developmental stages showing nucleus, mitochondrion, ER, Golgi apparatus, and apicoplast. ER rendering was made slightly transparent to allow better visibility of other features. Classifications were made based on cellular features typically associated with the respective developmental stages such as IMC cover or morphology of the parasite.

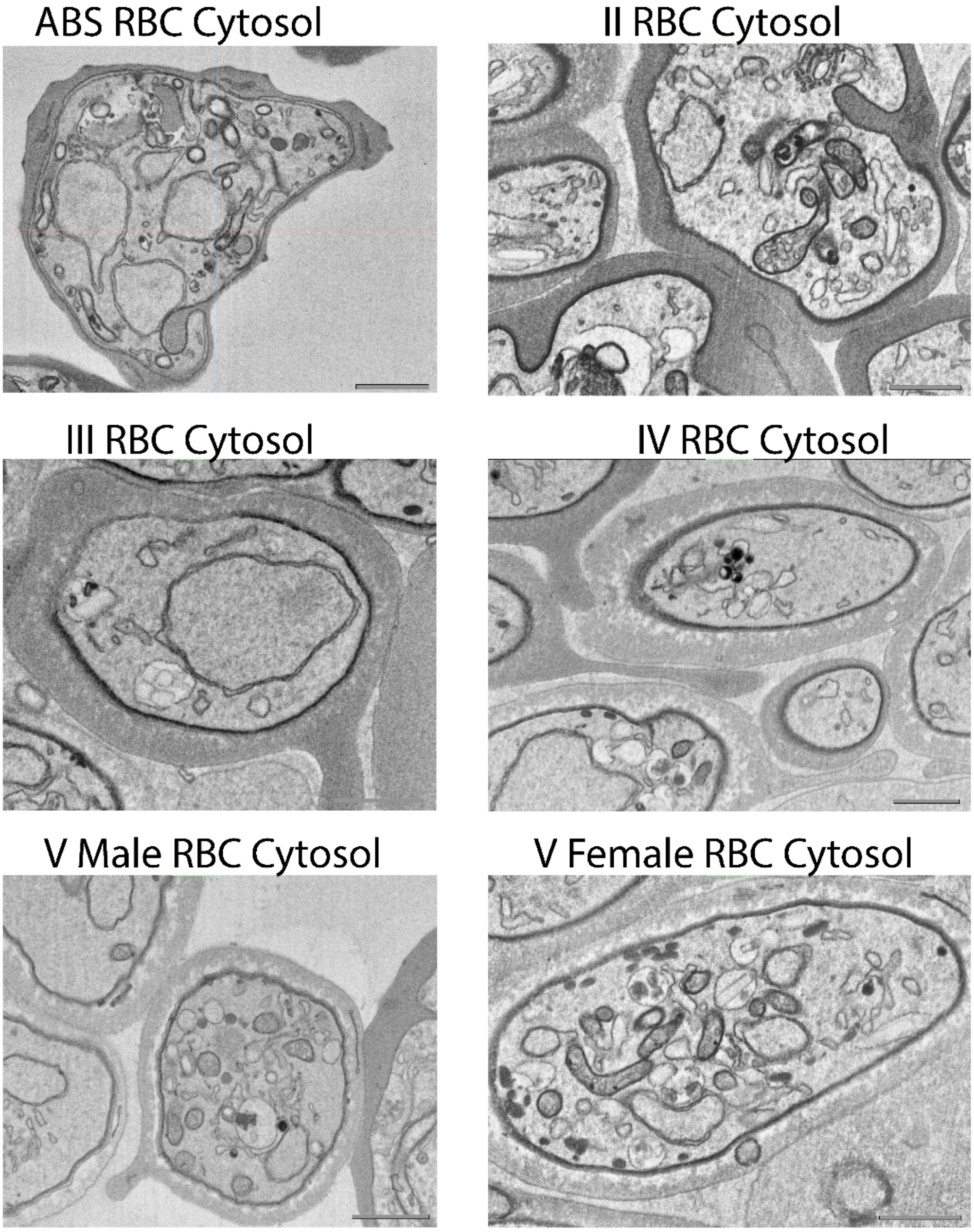


Figure S3. **Differing appearance of RBC cytoplasm in ABS and different gametocyte developmental stages.**
While RBCs infected with ABS and stage II gametocytes contain a homogenous and relatively more electron dense cytoplasm**.** Starting in stage III the cytoplasm becomes increasingly electron lucent and an even more electron lucent corona starts to develop around and in close proximity to the gametocyte.

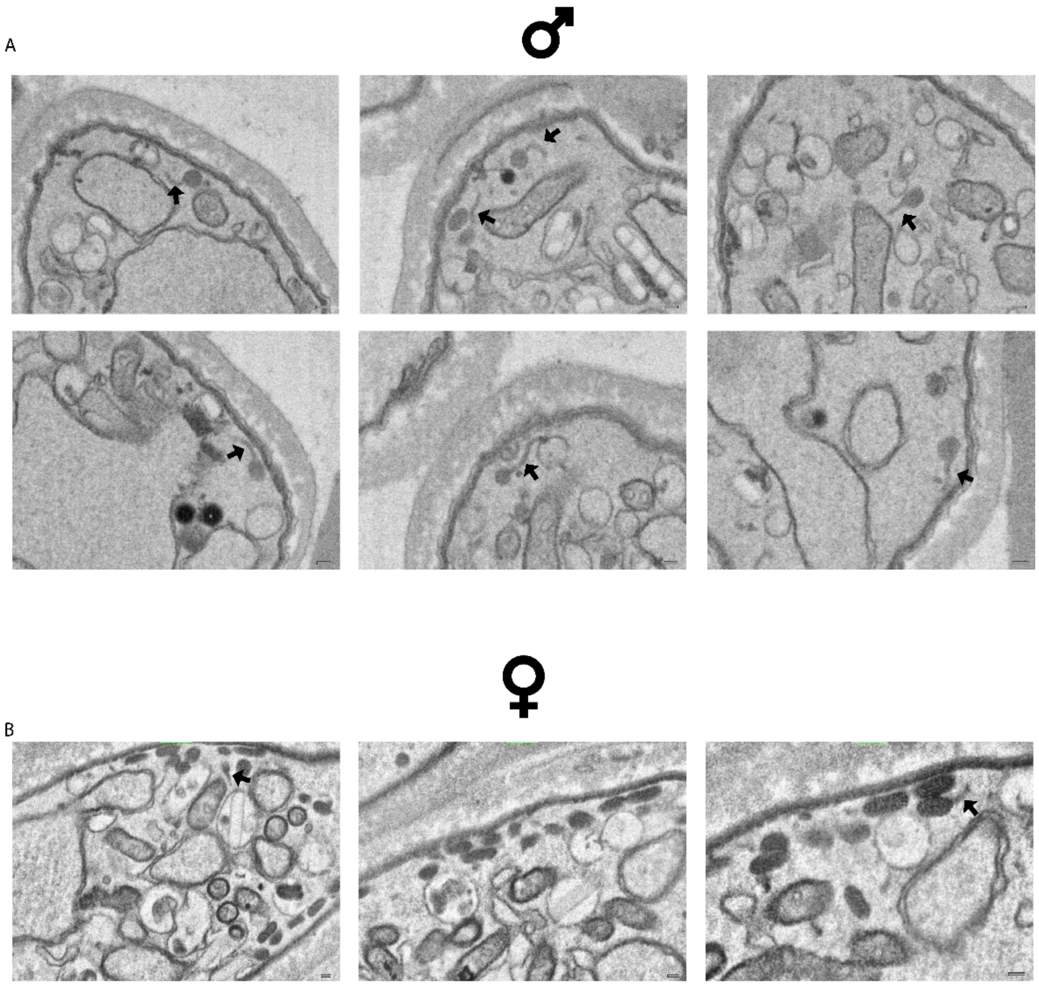


Figure S4. **Sex-specific differences in appearance of osmiophilic body-like structures.**
Male gametocytes contain vesicles that resemble osmiophilic bodies in their appearance, but with a tail-like appendix. These structures are only occasionally observed in female gametocytes. Exemplary micrographs showing typical OB morphology in (**A**) male and (**B**) female gametocytes. Note that most OBs observed in female gametocytes do not have tail extensions while the vast majority or all of the OBs observed in male gametocytes possess tail extensions. Arrows indicate tail extensions. Scale bars = 0.1 µm.

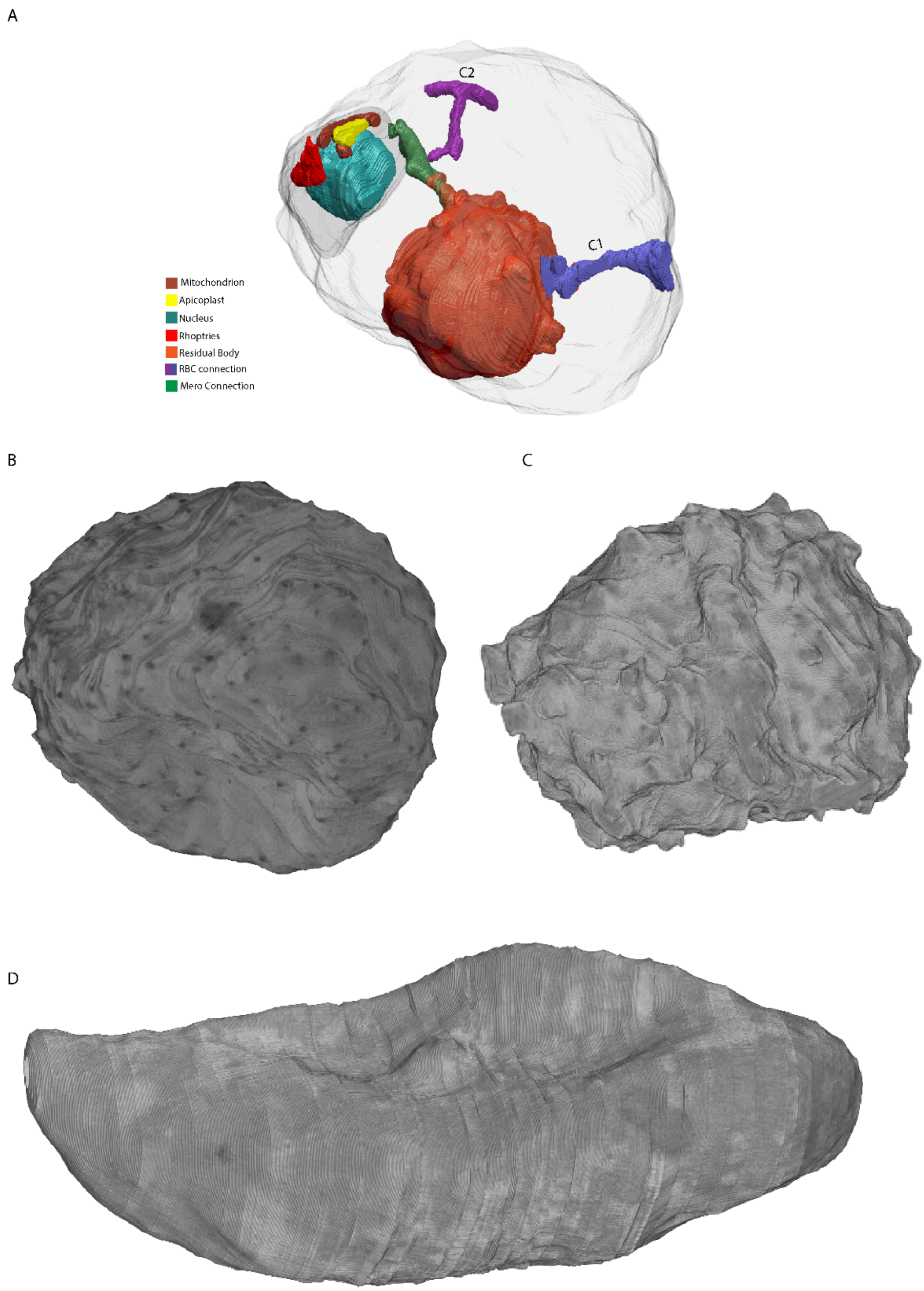


Figure S5. **Additional features of ABS.**
**A.** A segmented schizont harbors long tubular connections between the residual body and the RBC cytoplasm (C1, C2) and a daughter merozoite. **B-D.** Morphology of RBC infected by a developing schizont, a segmented schizonts, and a mature gametocyte with gray values from the underlying EM data.

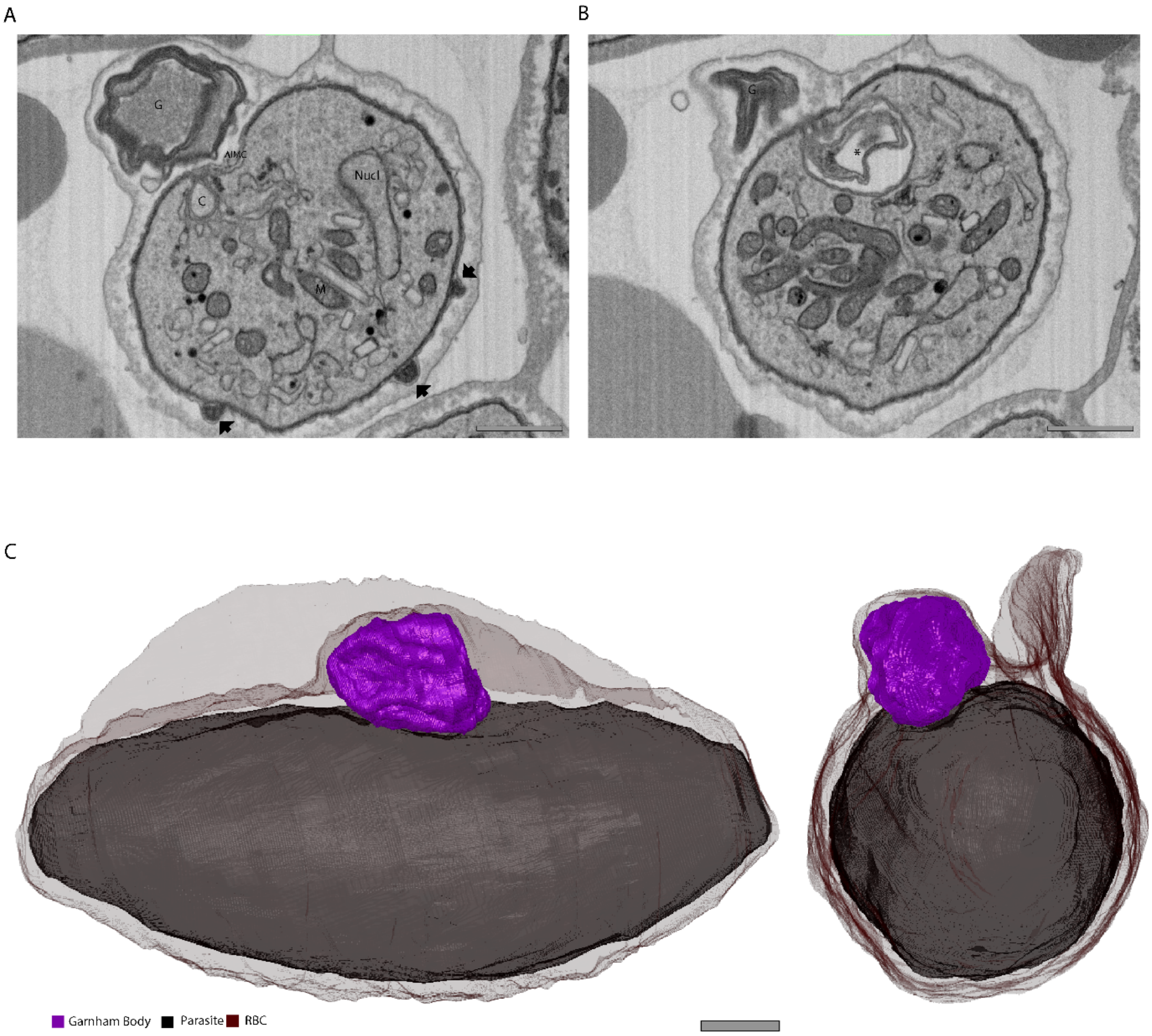


Figure S6. **The Garnham body.**
**A.** Exemplary micrograph showing membrane makeup of the Garnham body (G) and the presence of a cytostome (C), local lack of IMC (ΔIMC) and cellular protrusion (arrow). **B.** Exemplary micrograph showing aberrant internal structure opposing the Garnham body in the RBC (*). **C.** Rendering of the Garnham body from two angles. Scale bars = 1 µm.

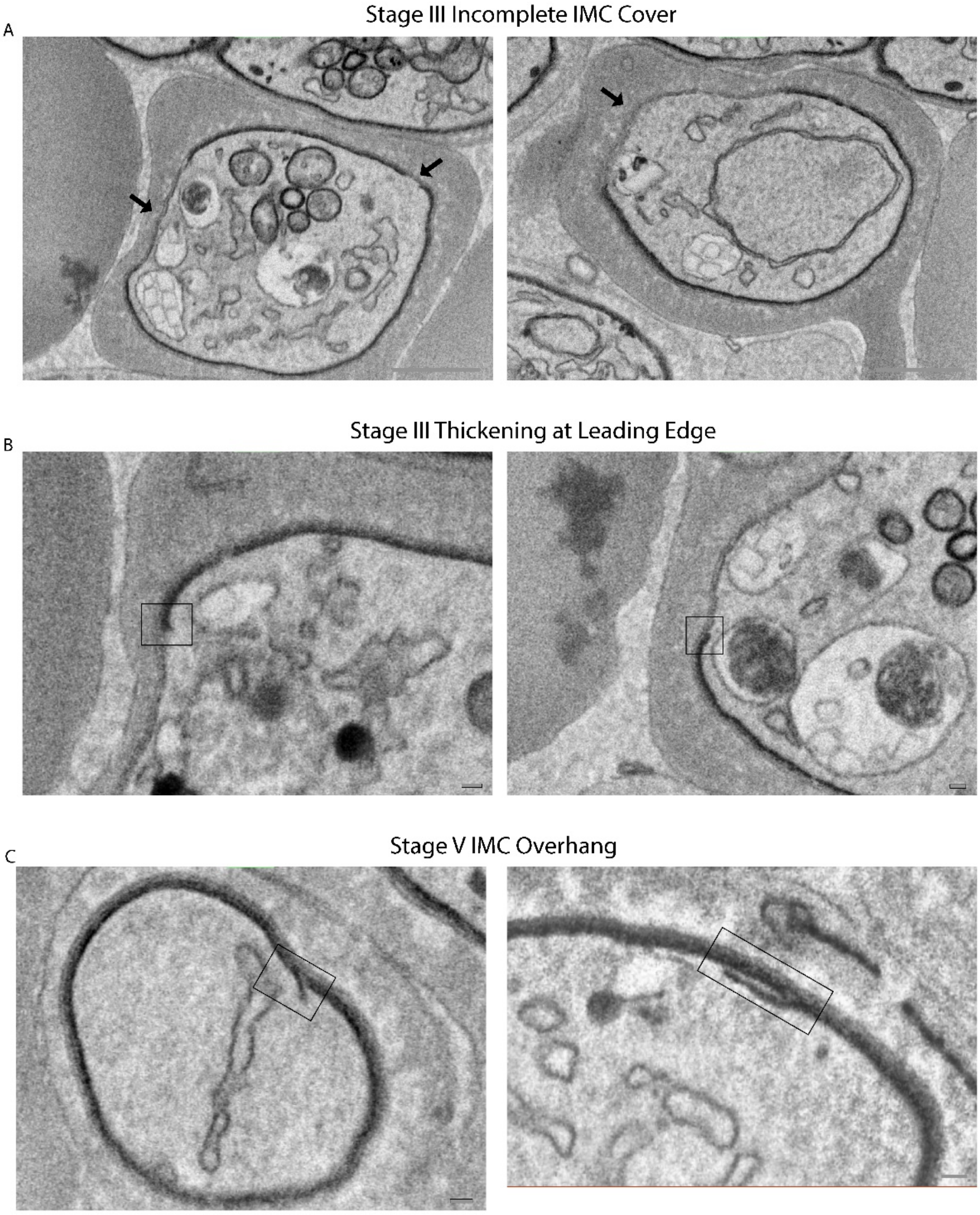


Figure S7. **IMC features in stage III and stage V gametocytes.**
**A.** Micrographs with examples of incomplete IMC cover in stage III gametocytes highlighted with arrows. Scale bars = 1 µm. **B.** Putative local thickening of IMC as described by Schneider *et al.* within boxes. Scale bars = 0.1 µm. **C.** IMC overhangs observed at polar ends of mature gametocytes. Scale bars = 0.1 µm.

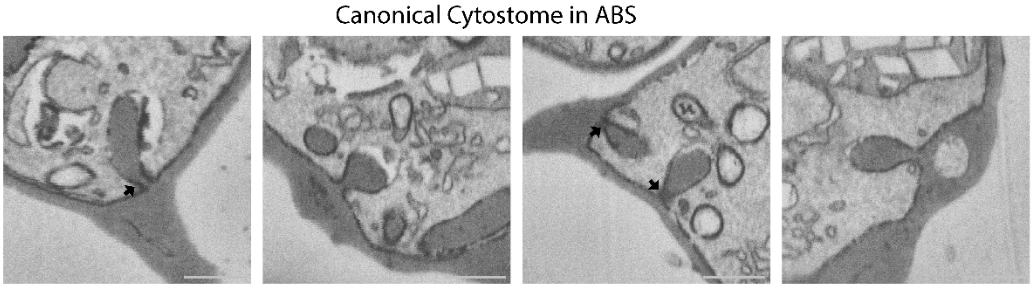


Figure S8. **The canonical cytostome.**
Example micrographs of the cytostome of trophozoites and schizonts. Arrows highlight the cytostomal collar. Scale bars = 1 µm.

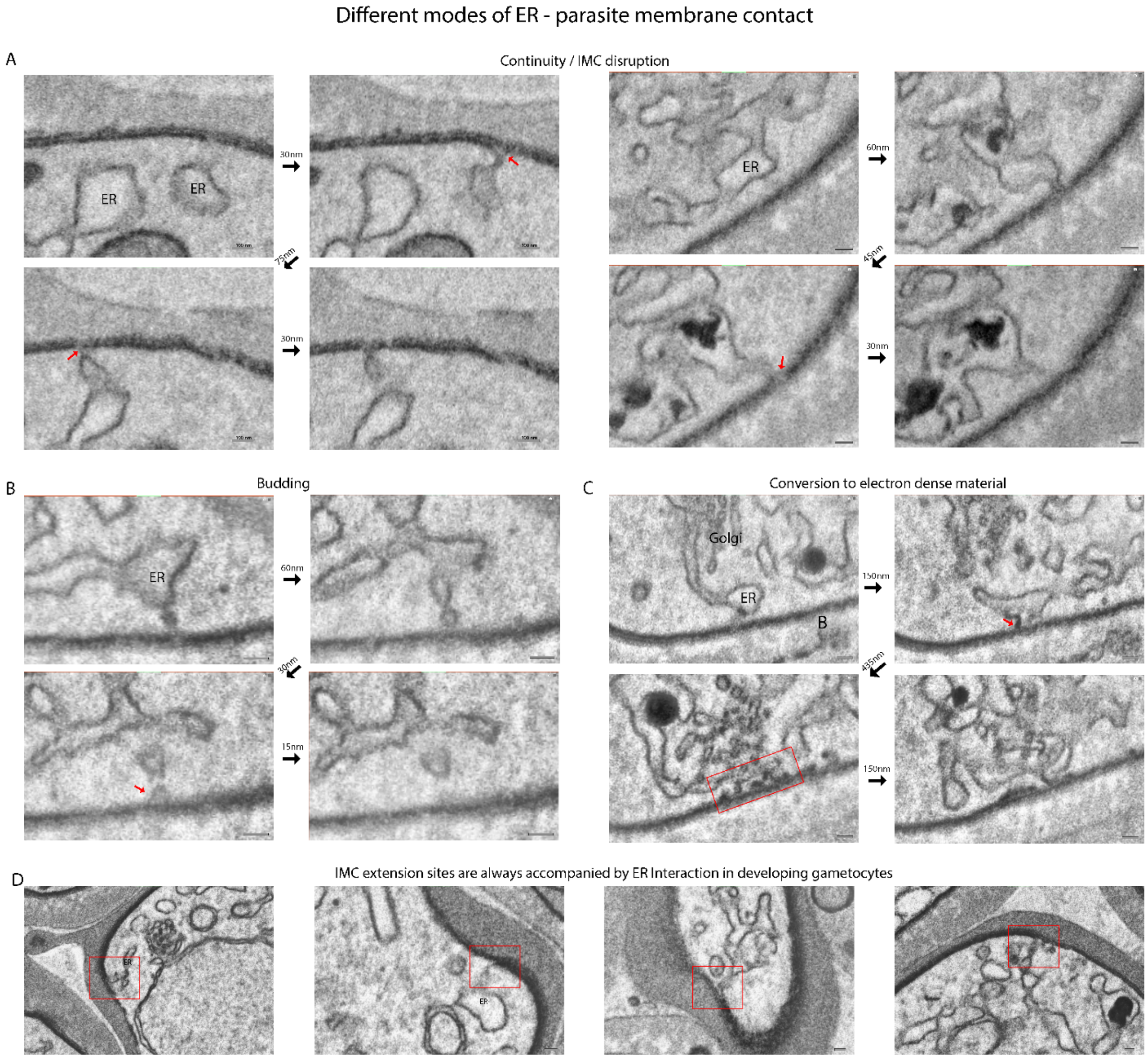


Figure S9. **Interactions between ER and parasite membrane.**

**A-C.** Series of micrographs from mature gametocytes showing ER in different putative interactions with the parasite membrane. **A.** The ER directly contacts and is continuous with the IMC, leading to a local disruption of the IMC. **B.** Same as (A) but connecting piece seemingly buds off from ER. **C.** ER is continuous with electron dense material that contacts the IMC. Black arrows between images express distance in the z-axis (acquisition plane) from one slice to the other. **D.** Exemplary micrographs showing contact of ER with the four IMC extension sites that could be identified in a developing gametocyte. All scale bars = 0.1 µm.

A

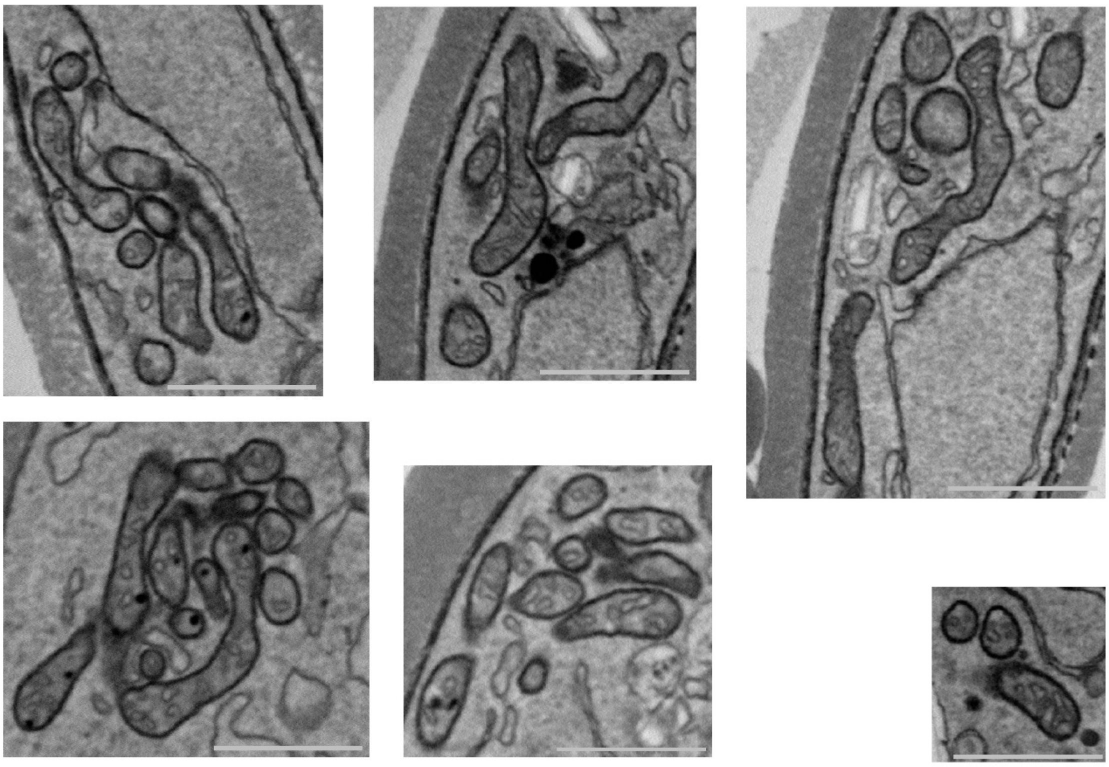

B

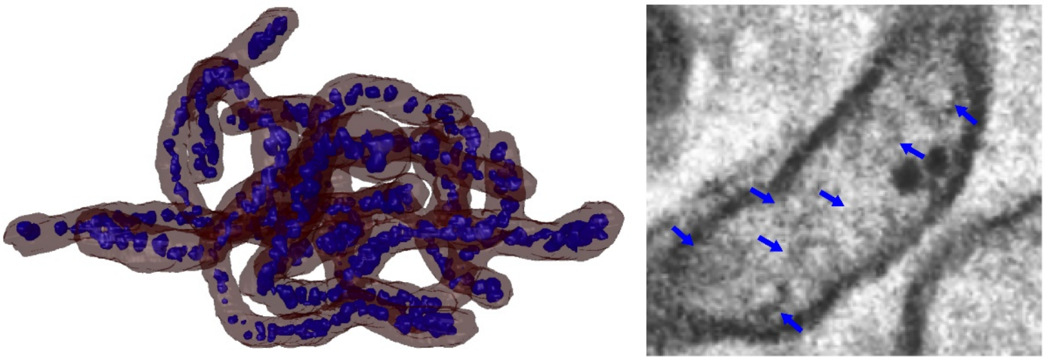

Figure S10. **Appearance and distribution of cristae in gametocytes.**
**A.** Exemplary micrographs from the low-noise gametocyte serial sectioning data. Note the presence of apparent branching points and proximity between individual cristae. The apicoplast often contained within a mitochondrial cluster is clearly distinct due the lack of internal membranous structures. Scale bars = 1 µm. **B.** Rendering of cristae (blue) within a gametocyte mitochondrion (red). Appearance of cristae in FIB-SEM data is highlighted in an exemplary micrograph through arrows.

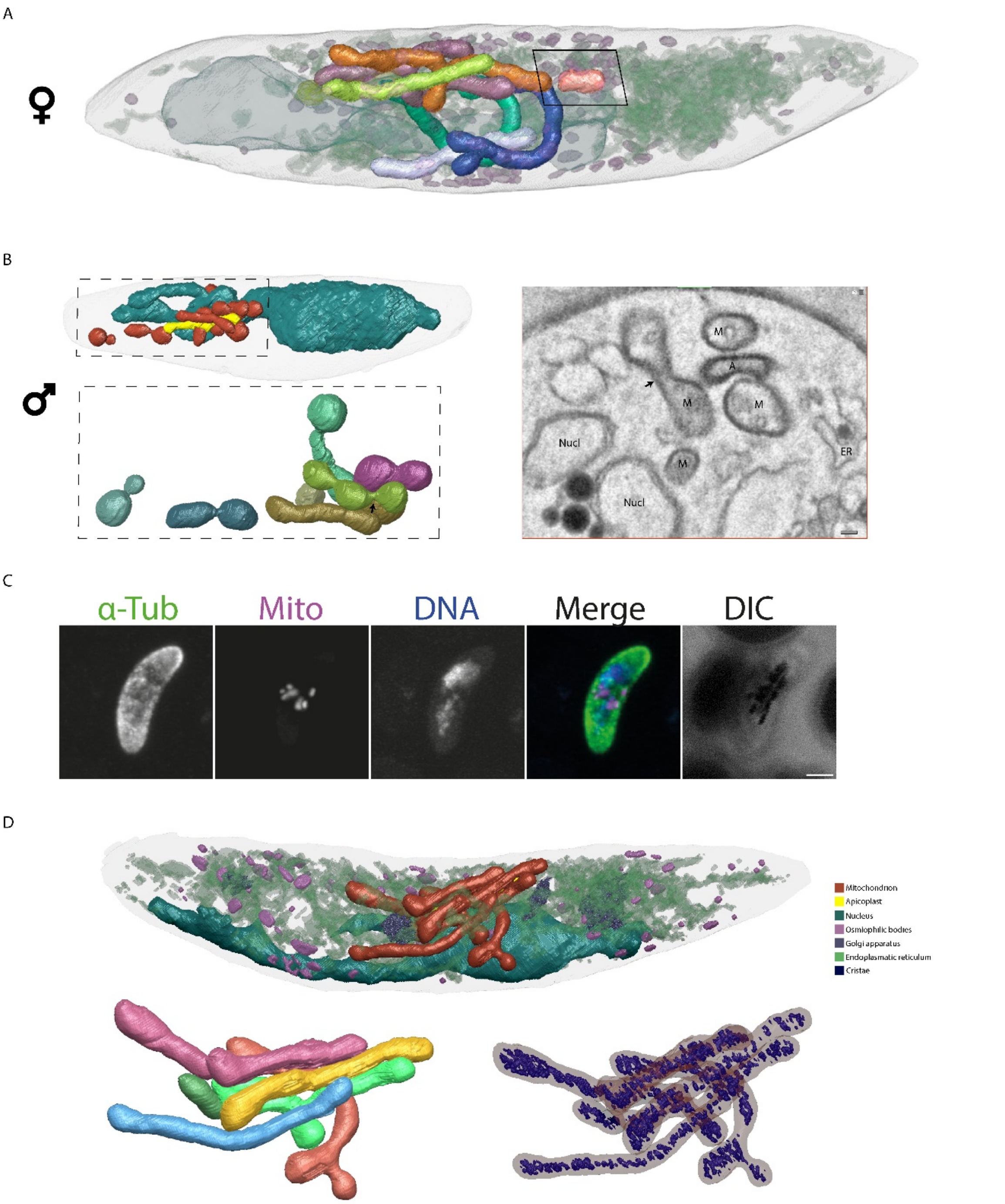


Figure S11. **Elaboration on multiple mitochondria in gametocytes.**
**A.** Rendering of a stage IV female gametocyte. Each of the seven mitochondria rendered in a separate color. One small mitochondrion is clearly recognizable as removed from the remainder (black rectangle). Nucleus, osmiophilic bodies, ER, and apicoplast are rendered with high transparency to provide cellular context. **B.** Rendering of nucleus, apicoplast, and dispersed mitochondria (top left), a zoom of only mitochondria without cellular context (bottom left), and an exemplary micrograph showing atypical mitochondrial morphology and constriction site (black arrow). **C.** Immunofluorescence analysis of a mature male gametocyte. Depicted left to right are maximum intensity projections using anti-α-tubulin antibodies, MitoTracker™, DAPI, merge of all channels and differential interference contrast (DIC) images. α-Tubulin was used to distinguish male and female gametocytes, DNA was visualized using DAPI, and mitochondria were visualized using MitoTracker. Mitochondrial staining suggests presence of multiple dispersed mitochondria. Scale bar = 2 µm. **D.** Renderings of a stage IV female gametocyte (upper section) and its mitochondrion highlighting the multiple mitochondria through different colors and distribution of cristae in a transparent mitochondrion (lower section).

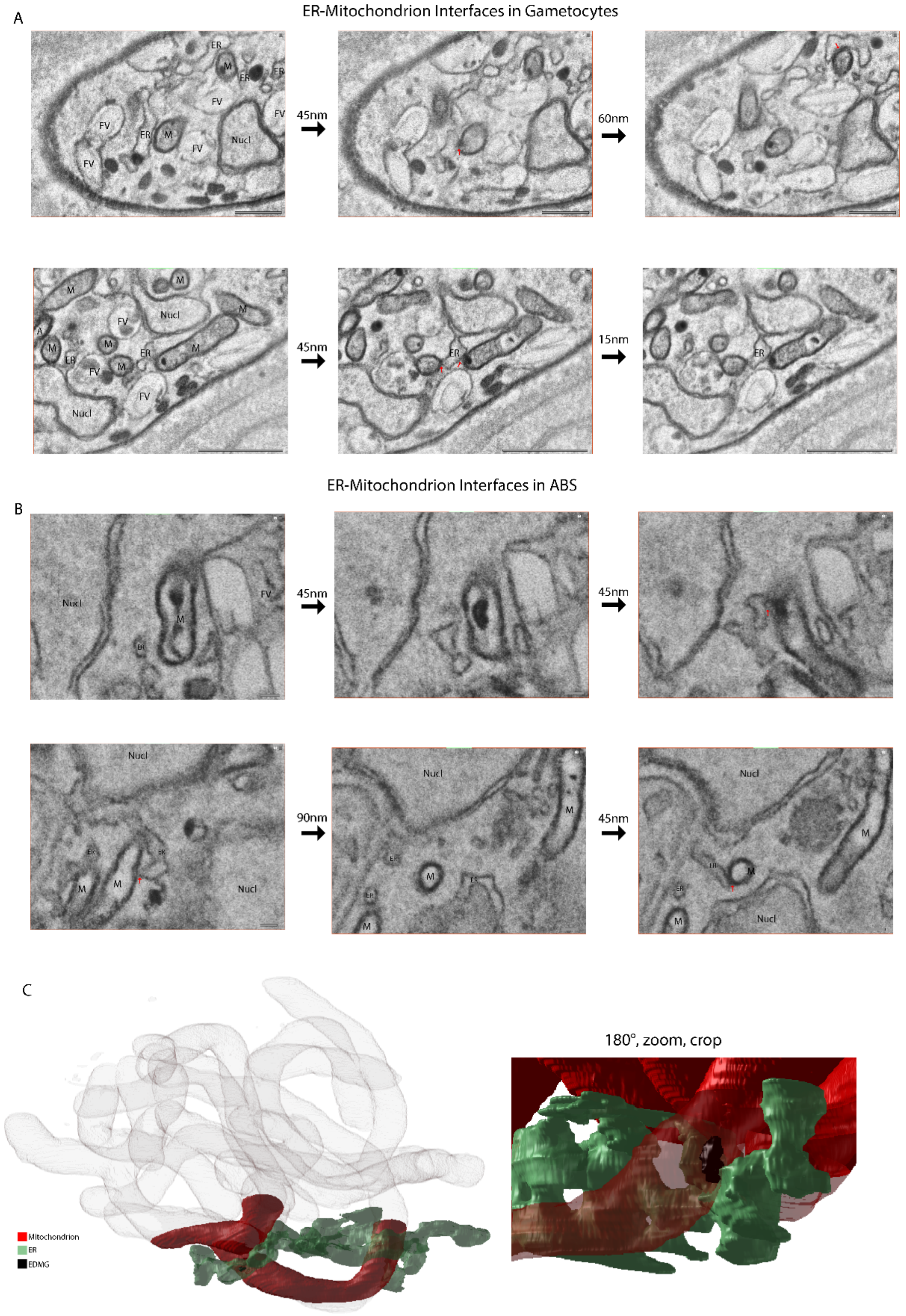


Figure S12. **ER-mitochondrion interfaces in gametocytes.**
Series of micrographs showing EDMGs spanning across mitochondrion and ER in **A**. gametocytes, scale bars upper panel = 0.5 µm, scale bars lower panel = 1 µm, and **B**. ABS, scale bars = 0.1 µm. **C.** Rendering of EDMG-ER-mitochondrion connection.

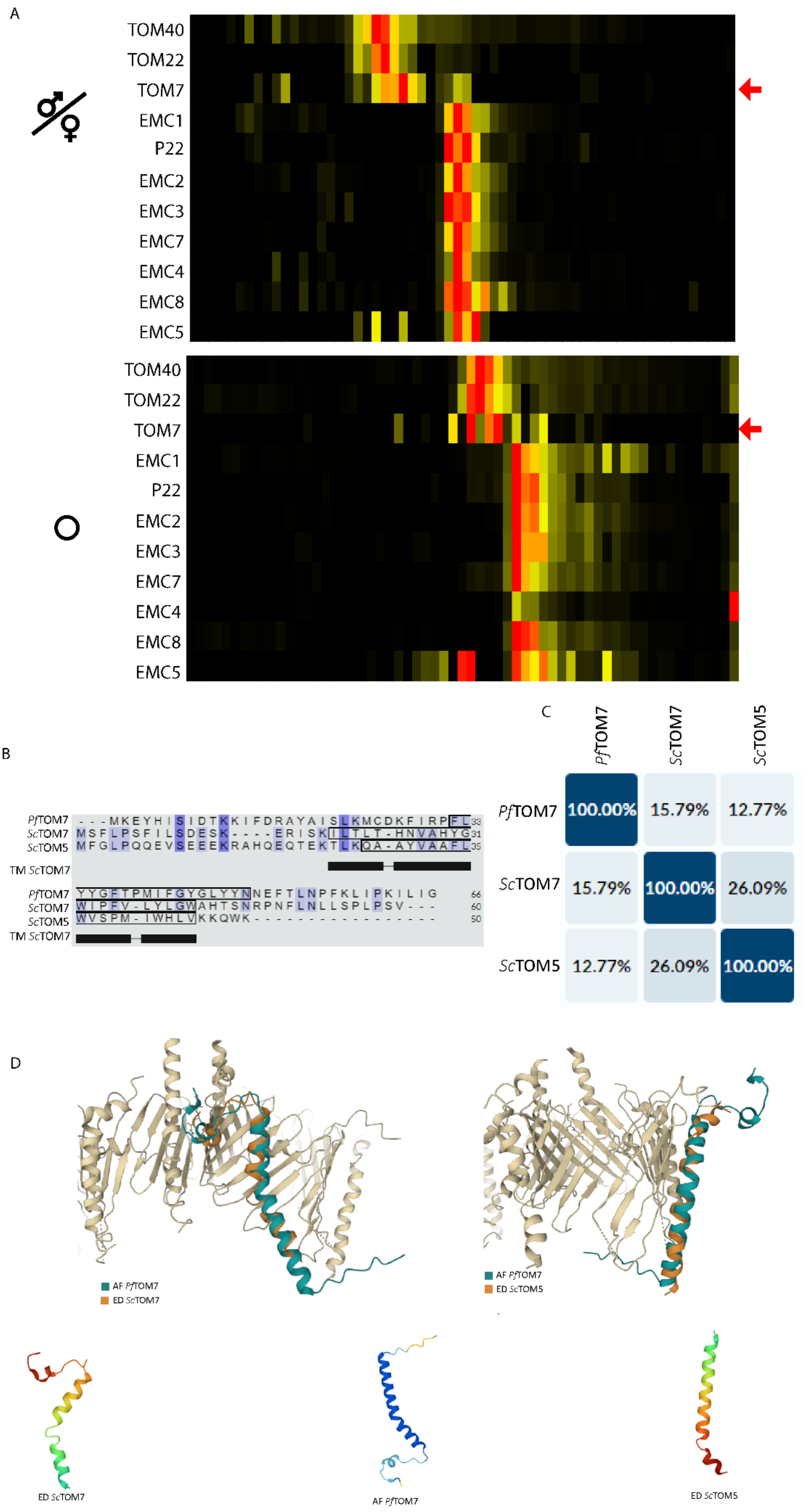


Figure S13. ***Pf*TOM7 as a putative EMC interactor and MCS facilitator.**

**A.** Heatmap showing migration of proteins (rows) in native gel electrophoresis (columns). High relative protein abundance is indicated. Appearance in the same column suggests overlapping migrations patterns and putative assignment to a protein interaction. Upper heatmap is derived from a gametocyte sample, lower heatmap is derived from an ABS sample. **B.** Multiple sequence alignment of *Pf*TOM7 with *Sc*TOM7 and *Sc*TOM5. Conserved residues have a purple background and residues with light-blue background are conserved in 2/3 sequences. Black boxes show presence of TM domains known from *Sc*TOM7. **C.** Sequence similarity matrix showing sequence identity between *Pf*TOM7, *Sc*TOM7, and *Sc*TOM5. **D.** Structural alignment of the *Pf*TOM7 AlphaFold prediction with the experimentally determined structures of *Sc*TOM7 and *Sc*TOM5 in the context of the experimentally determined structure of the TOM complex.
